# A conserved transcriptomic model defines metabolic resilience and vulnerability in obesity

**DOI:** 10.64898/2026.05.20.726524

**Authors:** Yin-Yuan Su, L Bundalian, Yin-Cheng Chen, E. Gjermeni, B. Gille, S. Richter, M. Jasaszwili, S. Palma-Vera, A. Hoffmann, Adhideb Ghosh, Christian Wolfrum, Matthias Blüher, S. Peleg, A. Garten, D Le Duc, Chen-Ching Lin

## Abstract

**Background:** Obesity arises from a complex interplay of genetic and environmental factors, with alterations of transcriptional networks that integrate metabolic, immune, and regulatory pathways. Conventional measures such as body mass index (BMI) quantify body size but fail to capture the molecular heterogeneity underlying divergent metabolic outcomes. We therefore sought to construct a gene expression-based transcriptomic representation of obesity, using BMI as a practical training anchor, and to use this framework to delineate transcriptional programs associated with metabolically healthy and pathogenic obesity, with subsequent projection to mouse transcriptomic data for cross-species validation.

**Methods:** Transcriptome data of human visceral adipose tissue (N= 1,298) were used to derive the transcriptomic BMI model, and genes contributing to the model were functionally annotated by gene set enrichment analysis. The human-trained model was subsequently applied to mouse selection lines (N = 18) with divergent obesity phenotypes. In the human cohort, post hoc stratification into metabolically healthy obesity (MHO) and metabolically unhealthy obesity (MUO) groups was performed using a downstream classification framework incorporating observed BMI together with predicted BMI, to assess whether model-derived predicted BMI reflected obesity-related pathophysiologic status.

**Results:** Model-selected genes were involved in coordinated regulation of lipid metabolism, immune activation, and growth signaling, extending to mitochondrial and translational pathways. Cross-species analyses uncovered conserved metabolic polarization: DU6 mice exhibited lipid-anabolic and inflammatory remodeling, whereas DU6P mice displayed oxidative, mitochondrial, and GH-axis–enriched transcriptional states. In human cohorts, MHO individuals showed upregulation of mitochondrial energetics and protein synthesis, while MUO individuals were characterized by increased autophagy, lipid catabolism, and stress-adaptive signaling on the transcriptional level. Together, these findings define a conserved molecular continuum linking oxidative efficiency to metabolic health and inflammation to metabolic vulnerability.

**Conclusions:** This integrative transcriptomic framework bridges human and mouse adipose biology to uncover conserved mechanisms underlying obesity phenotypes. By contrasting mitochondrial and translational programs with inflammatory and catabolic pathways, it provides mechanistic insight into metabolic resilience and a foundation for precision approaches to obesity management.

## Introduction

Obesity and its closely related cardiometabolic comorbidities—cardiovascular disease, type 2 diabetes mellitus (T2D), hypertension, and metabolic dysfunction associated liver disease — rank among the leading causes of morbidity and mortality worldwide^1^. Beyond these disorders, obesity substantially contributes to other major causes of death^2^, including cancer^3^, neurodegenerative diseases^4^, and chronic kidney disease^5^. However, obesity is not a biologically uniform condition. Individuals with excess adiposity can differ substantially in metabolic status, disease susceptibility and clinical course^6,7^.

Overweight and obesity are defined as an abnormal or excessive body fat accumulation that poses a health risk; a body mass index (BMI) ≥25 kg/m^2^ is considered overweight and ≥30 kg/m^2^ is considered obese^8^. However, BMI can both under- and overestimate adiposity and provides inadequate information about individual health risk^9,10^. Individuals with similar BMI, or even similar body-fat percentage, can exhibit markedly different cardiometabolic profiles^11,12^. The “personal fat threshold” hypothesis posits that the cardiometabolic impact of weight depends on each person’s capacity for safe fat storage, thereby explaining obesity-related heterogeneity, such as in T2D^13,14^. Nonetheless, BMI remains the dominant framework for defining obesity in both clinical and research settings^9,10^, underscoring the need for molecular approaches that better capture the underlying biology of adipose tissue and its contribution to metabolic health..

Murine models have been pivotal for discoveries in metabolism and aging, yet identifying systems that recapitulate human metabolic heterogeneity remains challenging. This challenge is compounded by the limited availability of cross-species transcriptional frameworks that directly connect human obesity heterogeneity with experimentally tractable animal biology^15,16^. The non-inbred Dummerstorf “Titan” mouse (DU6) is a particularly informative model of obesity^17^. DU6 was derived from a polygenic founder population (FZTDU) established in 1969/1970 by crossing four outbred with four inbred strains; selection for high body mass at six weeks began in 1975 with 60–80 mating pairs per generation, while an unselected control line was maintained throughout^18^. Adult DU6 mice average ∼110 g—substantially heavier than controls (∼44 g)—and accumulate ∼4% intra-abdominal fat versus ∼1% in controls. They also manifested most cardiometabolic disorders typical of metabolically unhealthy obesity (MUO)^19^. In contrast, the related Dummerstorf line DU6P, selected for high protein mass at six weeks, shows a “giant yet lean” phenotype with high muscle content^18,20^. These opposing phenotypes provide a tractable system to disentangle adipose tissue-driven lipid accumulation from lean-mass accrual.

To investigate whether obesity-related molecular states are conserved across species, we developed a transcriptome-based model trained on visceral adipose tissue gene-expression profiles from 1,298 human patients, using BMI as a practical training anchor. We focused on visceral adipose tissue (VAT) because this depot is closely linked to cardiometabolic disease, systemic metabolic regulation, and adipose inflammation, making it particularly well suited for interrogating molecular heterogeneity in obesity^22-24^. We then extended it across species by mapping high-confidence orthologous genes to adipose tissue samples from three well-characterized mouse lines: DU6 (obese-prone), DU6P (lean/protein-prone), and FZTDU (unselected control). This framework leverages adipose tissue transcriptomes to test whether conserved molecular programs differentiate metabolically unhealthy from healthier obesity states and to probe mechanisms underlying lipid accumulation versus oxidative efficiency.

## Methods

### Human data

The human cohort consists of 1,479 omental visceral adipose tissue samples sourced from the Leipzig Obesity BioBank (LOBB; https://www.helmholtz-munich.de/en/hi-mag/cohort/leipzig-obesity-bio-bank-lobb), which were collected in a clinical setting within a cross-sectional framework. The cohort included 432 men (BMI 49.12 ± 9.1 kg/m^2^; age 48.02 ± 11.8 years; 50.2% with T2D; 96.5% with BMI ≥ 30 kg/m^2^) and 1,047 women (BMI 48.47 ± 8.8 kg/m^2^; age 46.74 ± 11.8 years; 37.1% with T2D; 98.5% with BMI ≥ 30 kg/m^2^). Participants underwent comprehensive clinical phenotyping with respect to obesity status, metabolic parameters, and anthropometric characteristics^25,26^. Adipose tissue samples were obtained during planned laparoscopic abdominal procedures, as previously reported^27^. Individuals were excluded if they were younger than 18 years, had a history of chronic drug or alcohol abuse, had smoked within the year prior to surgery, or presented with acute inflammatory conditions, advanced malignant disease, uncontrolled thyroid dysfunction, or Cushing’s syndrome.

### Library preparation and RNAseq analysis of human data

Total RNA was extracted from adipose tissue samples following the SMART-seq protocol (Switch Mechanisms at the 5’ End of RNA Templates)^28^. Sequencing libraries were then constructed and subjected to single-end sequencing on a NovaSeq 6000 instrument at the Functional Genomics Centre Zurich.

Raw sequencing reads were processed with Fastp (v0.23.4) to remove adapters and perform quality trimming^29^. The cleaned reads were aligned to the human reference genome (GRCh38, GENCODE release 47) using STAR (v2.7.11b), permitting up to 50 multiple alignments per read^30^. Gene-level quantification was carried out with featureCounts (v1.5.3), applying fractional assignment for reads mapping to multiple genomic locations^31^.

### Origin of mouse strains and housing conditions

The mouse lines included in this study stem from a long-term selection experiment, which has been conducted in the Research Institute for Farm Animal Biology, generating, among others, mouse lines with increased body mass (DU6)^32^ and protein mass (DU6P)^17^. These mice have been maintained for more than 140 generations alongside an unselected control line (FZTDU)^33,34^. All animal procedures were performed in accordance with national and international guidelines and ethically approved by our own institutional board (Animal Protection Board from the Leibniz Institute for Farm Animal Biology). Mice were housed in a specific pathogen-free (SPF) environment with defined hygienic conditions at a temperature of 22.5 ± 0.2 °C, at least 40% humidity and a controlled light regime with a 12:12 h light-dark cycle. The mice were kept in polysulfone-cages of 365 × 207 × 140 mm (H-Temp PSU, Type II L, Eurostandard, Tecniplast, Germany) and had free access to pellet feed and water. Mice were fed ad libitum using a SBF (gross energy (GE) = 16.7 MJ kg^−1^, metabolizable energy (ME) = 14.0 MJ kg^−1^, Supplementary Table 2) including 12% fat, 27% protein, and 61% carbohydrates (ssniff® M-Z autoclavable, Soest, V1124-300, Germany) ^17^.

Male mice were sacrificed at 8–9 weeks of age, and inguinal and epigonadal white adipose tissue was resected and either immediately frozen for RNA isolation or fixed in 4% paraformaldehyde for histological analyses.

### Adipose tissue histology and immunohistochemistry

Paraffin-embedded white adipose tissue (WAT) was sectioned into 10 µm slices, dewaxed, dehydrated in an alcohol series, and stained with hematoxylin (Carl Roth, T865.1) and eosin (Carl Roth, 3137.1).

For immunohistochemical detection of F4/80 as macrophage marker a rat monoclonal antibody anti-mouse F4/80 was used (Cat.-No: 123101, Biolegend, 1:400 dilution). For immunohistochemical detection of macrophages, tissue sections were deparaffinized, and quenched with 3% hydrogen peroxide for 10 min. Slides were heat retrieved with citrate buffer at pH 6.1 (Catalog S1699, DakoCytomation) for 4 min at 123°C, then cooled for 15 min, followed by a running tap water rinse. Sections were then incubated with the anti-F4/80 antibody at room temperature for 60 min and rinsed with TBS. F4/80 stained sections were incubated for 30 min in Rat Probe and then Rat Polymer-HRP (Rat-on-Mouse HRP Kit, Cat-No: RT517H, Biocare Medical/Zytomed systems). Slides were rinsed with TBS and developed with DAB-Plus (DakoCytomation), counterstained in modified Mayer’s hematoxylin and blued in 0.3% ammonia water, followed by a tap water rinse^35^.

Brightfield images were taken with an Keyence BZ-X (Keyence Corporation of America) at 20 x magnification (for adipocyte size determination: FTZDU n = 9 epigonadal, n = 7 inguinal, DU6 n = 12 both inguinal and epigonadal, DU6P n = 6 both inguinal and epigonadal; for evaluating F4/80 staining/crown-like structures: FTZDU n = 6 for epigonadal and n = 4 inguinal, DU6 n = 12 for both epigonadal and inguinal, DU6P n = 4 epigonadal and n = 1 inguinal). For adipocyte size, three images per sample and for crown-like structures, at least two images per sample were analysed. Adipocyte size was measured using the ImageJ^36^ plugin Adiposoft^37^, crown-like structures were counted manually and normalized to adipocyte number as determined by Adiposoft analysis. Statistically significant differences between 3 groups were tested with one way Brown-Forsythe and Welch ANOVA followed by Dunnett T3 correction for multiple comparisons and between two groups were tested with Mann-Whitney non-parametric test and visualized using GraphPad Prism 10.

### Data Source and RNA Sequencing of Dummerstorf Mouse Lines

Tissue samples were collected and isolated from the three Dummerstorf (DU)/Titan mouse lines a) FZTDU (normal), DU6 (obese) and DU6P (lean) each having 6 biological replicates. Library preparation was done using the Illumina RNA preparation kit following the protocol instructed by the manufacturer. Raw sequencing data was preprocessed using the nf-core/rnaseq pipeline. The quality was evaluated using *FastQC* and adapter sequences along with the low-quality bases were trimmed using *cutadapt*. Processed reads were aligned to the *Mus musculus* (GRCm39) reference genome using STAR while the quantification is executed using HTseq.

### Data preprocessing

To investigate pathways and functions related to human obesity mechanisms in mice, we trained a BMI prediction model and applied it to mouse data. Protein-coding genes were extracted from the *Mus musculus* genome assembly (GRCm39) and the *Homo sapiens* genome assembly (GRCh38.p14). Genes with zero counts across all samples within a species were removed, yielding 19,884 human genes and 19,087 mouse genes for analysis.

Starting from the Kallisto-derived expression estimates, we applied quantile normalization separately to 1,479 human visceral adipose tissue samples and 18 mouse epigonadal adipose tissue samples (6 DU6, 6 DU6P, 6 FZTDU) to minimize expression-level perturbations and ensure comparable within-species distributions. Subsequently, feature-wise Z-score standardization was performed within each species to place all genes on a common scale, thereby reducing biases in cross-species prediction arising from intrinsic biological variation.

High-confidence orthologous gene pairs between human and mouse were retrieved from Ensembl Compara^38,39^. Orthologs were considered high confidence when supported by concordant gene-tree/species-tree reconciliation, sufficient sequence similarity and alignment coverage, conserved gene structure, and synteny across species. After orthology mapping, 16,042 high-confidence human–mouse orthologs were retained as features for subsequent BMI model development.

### Application of the human-trained BMI-anchored model to mouse transcriptomes

To focus the analysis on obesity, we restricted the training set to individuals with BMI ≥ 30. Because extremely high BMI (morbid obesity) may reflect mechanistically distinct regimes of fat accumulation and metabolism, we further excluded the top 10% of BMI values to mitigate distributional shift and focus on the predominant cohort (Supplementary Fig. S1). The resulting training set comprised 1,298 human visceral adipose samples with BMI 30–60.

To obtain a sparse, transfer-ready signature, we trained a LASSO (L1-regularized) linear regression model to predict human BMI from gene expression. Given the multifactorial nature of BMI and potential non-linearities at the global scale, model selection emphasized rank consistency, evaluating performance with Spearman’s rank correlation (ρ) in addition to mean squared error (MSE). The regularization strength was tuned to achieve a prediction Spearman’s ρ of approximately 0.8, thereby balancing parsimony and stability over marginal improvements in in-sample fit.

After training, we applied the learned human coefficients to the expression of the corresponding mouse orthologs, restricting to features with non-zero coefficients in the human model. For each mouse sample, the orthologous expression vector was aligned to the human feature set, and the linear predictor was used to generate mouse BMI predictions. Line-level summaries were then examined across the three mouse lines (DU6, DU6P, FZTDU) for downstream analyses.

### Identification of functional enrichment within model genes and group differences

To elucidate the biological processes represented by model-selected genes and group-specific differences, we performed gene set enrichment analysis (GSEA)^40,41^. This approach was used to identify biological functions enriched in association with BMI prediction as well as in group contrasts (e.g., DU6 vs. DU6P and MUO vs. MHO). Therefore, GSEA was applied to both model–selected genes and group-difference gene rankings, using Gene Ontology (GO) terms with a hierarchical depth greater than five to ensure adequate functional specificity.

For model-based analyses, human genes selected by the model were ranked according to their regression coefficients. For group-difference analyses, genes were ranked using log2 fold changes or corresponding expression-derived effect sizes, depending on data availability.

For differential expression analysis, in the mouse dataset, where raw read counts were available, differential expression analysis was performed using DESeq2^41^, and genes were ranked by the resulting log2 fold changes. However, in the human dataset, where only Kallisto-derived expression values were available, group differences were quantified using expression-derived log2 fold-change– like effect sizes. Specifically, for each gene in each sample, expression was first scaled by the geometric mean of that gene within the corresponding group and then normalized by the within-sample median. Group-level effect sizes were subsequently calculated as the log2 ratio of the mean normalized expression values between groups. These human-derived effect sizes were used solely as the ranking metric for downstream GSEA and were not involved in model training.

For each analysis, an enrichment score (ES) was computed to quantify the degree to which a GO term was overrepresented within the ranked list. Statistical significance was assessed using 1,000 permutation tests, and ES values were standardized into z-scores as normalized enrichment score (NES). Functional modules with z ≥ 2 were considered significantly enriched in the corresponding gene sets.

### Patient stratification using LOWESS and decile standardization

To address systematic bias in predicted versus observed BMI, we employed locally weighted scatterplot smoothing (LOWESS) regression as a calibration approach. Predicted BMI values were treated as the independent variable (x), and observed BMI values as the dependent variable (y). A LOWESS curve was fitted using parameters frac = 0.3, it = 3, and degree = 1, which balances local flexibility with global smoothness while reducing the influence of outliers. For each patient, we calculated the residual relative to the fitted curve:

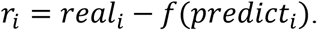

To account for heteroscedasticity, the conditional variance of residuals was estimated by applying LOWESS to the squared residuals, yielding local standard deviations *local*_*std*(*predict*_*i*_). Each residual was standardized as:

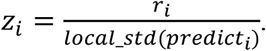

Because even standardized residuals may show systematic dependence on real BMI, we performed an additional decile-based normalization. Patients were divided into deciles according to their observed BMI. Within each decile d, the mean *µ*_*d*_ and standard deviation *σ*_*d*_ of *z*_*i*_ were calculated, and each patient’s residual was further standardized:

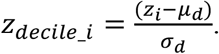

*T*his procedure ensures that patients are evaluated relative to peers with similar BMI, preventing systematic misclassification of low-BMI patients as MUO and high-BMI patients as MHO. Final group assignments were based on these decile-standardized scores:

- MHO: *z*_*decile_i*_ ≥ 1
- MUO: *z*_*decile_i*_ *≤* -1
- Neutral: -1 < z_decile_i < 1

This decile standardization framework allows patient stratification to reflect true molecular-to-phenotypic discordance, rather than statistical artifacts of regression to the mean.

### Calculation for the contiguous run/stretch of homozygosity scores (ROH)

Genomic DNA of DU6, DU6P and FZTDU control mice was sequenced by whole genome sequencing (WGS) analysis as described in (PMID: 35189878).

Runs of homozygosity (ROH) were identified as continuous genomic segments containing predominantly homozygous genotypes. For each candidate stretch, we quantified the degree of homozygosity by computing a homozygosity score^42^ defined as:

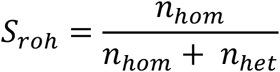

where n_hom_ and n_het_ denote the numbers of homozygous and heterozygous single-nucleotide polymorphisms (SNPs), respectively, within the segment (excluding missing genotypes). The scores were further normalized to account for their gene length i.e. dividing the S_roh_ by their corresponding gene lengths. This score reflects the individual’s overall level of auto zygosity, with higher values indicating increased genomic homozygosity and potential recent parental relatedness. The threshold was defined by calculating the 90th percentile of the normalized scores to identify outlier homozygosity regions indicative of possible auto zygosity^43^.

## Results

### Construction of a BMI-anchored transcriptomic model and functional annotation of model-associated genes

To investigate molecular mechanisms underlying obesity progression, we established a transcriptome-based model to derive predicted BMI from visceral adipose tissue RNA-sequencing profiles. RNA-sequencing data from 1,298 human visceral adipose tissue samples with BMI 30-60 were used as input features, with measured BMI serving as the training label. BMI was selected because it is routinely collected and clinically relevant, whereas other metabolic parameters contained substantial missing data and were therefore less suitable for model training. Rather than optimizing the model solely for maximal prediction accuracy, we tuned the model to achieve a Spearman’s rank correlation of 0.8 between predicted and measured BMI, thereby preserving BMI-related ordering while avoiding excessive dependence on the observed phenotype. This calibrated framework enabled predicted BMI to serve as a transcriptomic representation of obesity-related biological status for downstream analyses.

To evaluate the biological relevance of model-selected genes, we performed gene set enrichment analysis (GSEA) to identify enriched Gene Ontology (GO) terms, which were subsequently grouped into functional themes (Supplementary Methods). Normalized enrichment scores (NES) revealed contributions from diverse cellular processes (Fig. 1).

**Figure 1.**
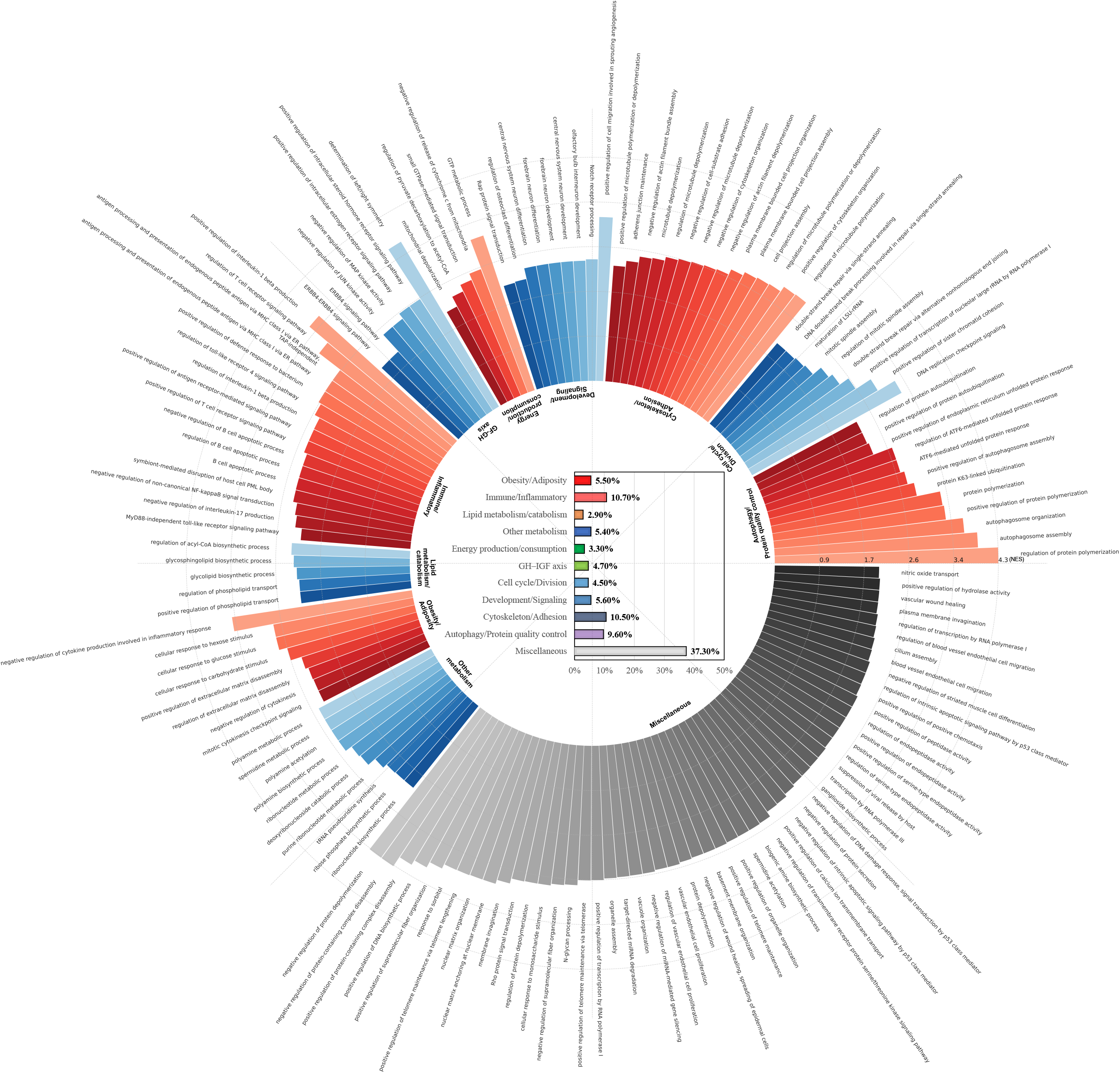
Circular representation of relative enrichment contributions of individual functions grouped by biological category. Circular bar plot depicting the normalized enrichment scores (NES) of individual functions identified from Gene Set Enrichment Analysis (GSEA) of model-selected genes. Each bar represents a single biological function, with bar length corresponding to its NES value. Functions are grouped into higher-order biological categories, labeled in the inner circle. The color-coded inner bar plot indicates the relative enrichment contribution of each category, calculated as the proportional sum of NES values of functions within that category relative to the total NES. This visualization highlights the integrated transcriptional architecture of obesity, demonstrating coordinated contributions from metabolic, immune, structural, and regulatory processes.

Functions directly related to obesity were prominent, encompassing both the obesity/adiposity (5.5%) and immune/inflammatory (10.7%) categories. Enriched processes included cytokine regulation, adipose tissue inflammation, and glucose-responsive pathways, underscoring the close interplay between immune signaling and adipose metabolic regulation. Broader metabolic programs were also enriched, notably lipid metabolism/catabolism (2.9%) and other metabolism (5.4%), which involved lipid-handling, polyamine, and nucleotide-associated processes, indicating that both lipid and non-lipid metabolic pathways are represented in the model-selected genes. A third major theme corresponded to growth and tissue regulation, comprising energy production/consumption (3.3%), GH–IGF axis (4.7%), cell cycle/division (4.5%), development/signaling (5.6%), and cytoskeleton/adhesion (10.5%). These categories collectively represent cellular energetics, hormone signaling, proliferation, and structural remodeling. Additionally, autophagy/protein quality control (9.6%) fraction captured diverse regulatory processes.

In summary, these findings demonstrate that BMI-associated transcriptional programs extend beyond metabolism to engage immune, metabolic, and structural pathways that collectively shape adipose tissue growth and function.

### Application of the model to mouse lines with divergent obesity phenotypes

To assess the cross-species relevance of our model, we applied it to transcriptomic data of epigonadal adipose tissue from three well-characterized mouse selection lines: DU6, selected for high body weight; DU6P, selected for high protein mass; and FZTDU, the unselected control. The condition factor (CF) ranked DU6 *≈* DU6P > FZTDU (Fig. 2a). In contrast, the relative content of white adipose tissue followed the order DU6 > DU6P *≈* FZTDU (Fig. 2a), revealing a discrepancy between gross phenotype (CF) and adipose tissue composition. Consistent with this divergence, independent histological analyses performed in a separate cohort showed adipose morphological differences among the three mouse lines, including adipocyte hypertrophy and inflammatory remodeling in DU6 (Supplementary Fig. S2).

**Figure 2.**
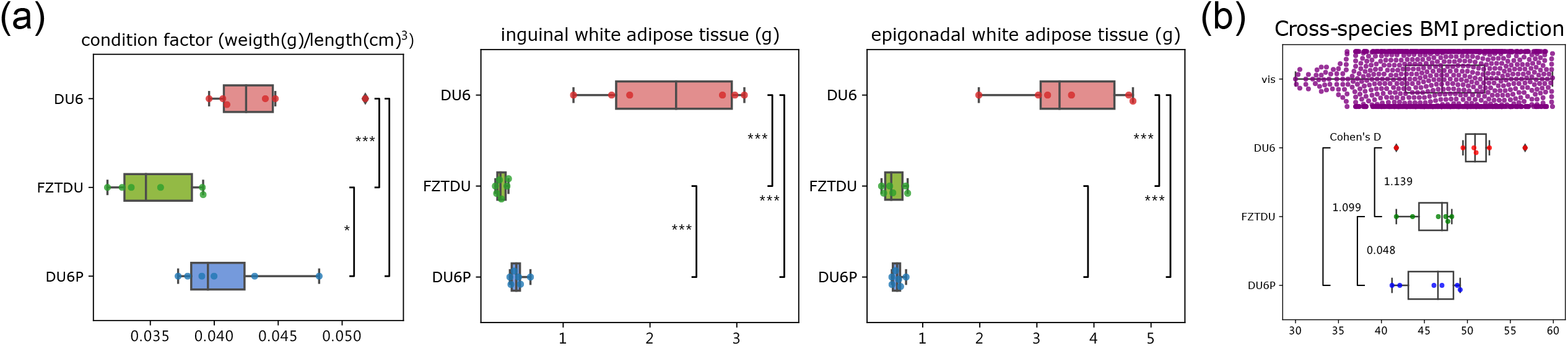
Phenotypic and molecular divergence among Dummerstorf (Titan) mouse lines. (a) Boxplots showing the distribution of the condition factor (weight [g]/length [cm]^3^) and normalized white adipose tissue (WAT) weights, calculated as depot weight divided by total body weight, from the inguinal and epigonadal depots across the Dummerstorf mouse lines (DU6, DU6P, and FZTDU). Statistical significance of pairwise group differences was evaluated using the two-sided Mann– Whitney U test; significance thresholds are denoted as * p < 0.1, ** p < 0.05, *** p < 0.01, and blank comparisons indicate non-significant differences. (b) Cross-species BMI prediction based on the human visceral adipose tissue transcriptome model. Each point represents an individual sample: human visceral adipose tissue samples (BMI 30–60, n = 1,298) and adipose tissue from the three Dummerstorf lines (DU6, DU6P, and FZTDU). Boxplots summarize group distributions, and effect sizes (Cohen’s d) are shown for pairwise comparisons. The predicted BMI ranking (DU6 > DU6P *≈* FZTDU) mirrors adipose-tissue accumulation rather than external body size, while aligning with the human visceral reference distribution. Together, the plots highlight a decoupling between gross phenotype and adipose tissue molecular signatures of adiposity.

When the human-trained BMI-anchored transcriptomic model was applied to mouse adipose transcriptomes, the resulting predicted BMI values ranked DU6 > DU6P *≈* FZTDU (Fig. 2b), mirroring the order of epigonadal white adipose content rather than CF. This result indicates that the model captures molecular signatures of adiposity that are not reflected in external body measures alone. Thus, the the BMI-anchored transcriptomic model distinguishes between animals with similar condition factors but divergent fat composition, underscoring its ability to extract biologically meaningful signals of obesity from transcriptomic profiles.

### Opposing lipid-anabolic and oxidative-catabolic states define DU6 and DU6P along the predicted-BMI axis

To determine whether the BMI-trained model captures biologically meaningful distinctions in obesity states, we first performed Gene Set Enrichment Analysis (GSEA) to assess broad functional trends between DU6 (obese-prone) and DU6P (lean/protein-prone) mice.

Enrichment profiles revealed contrasting transcriptional programs between the two lines (Fig. 3a, Supplementary Fig. S3, and S4). Relative to DU6P, DU6 showed stronger enrichment of immune/inflammatory pathways, including cytokine regulation, leukocyte migration, and mast cell degranulation, alongside broad activation of cell cycle/division processes, consistent with comparatively greater inflammatory activation and proliferative remodeling. By contrast, DU6P showed relative enrichment of growth and tissue regulation, dominated by energy production/consumption and development/signaling processes. Additional enrichment of lipid metabolism/catabolism and other metabolism in DU6P suggested metabolic flexibility, whereas immune/inflammatory and cell cycle/division were notably absent. Together, these findings indicate a functional divergence between the two lines, with DU6 biased toward immune and proliferative programs and DU6P biased toward energetics- and development-associated programs.

**Figure 3.**
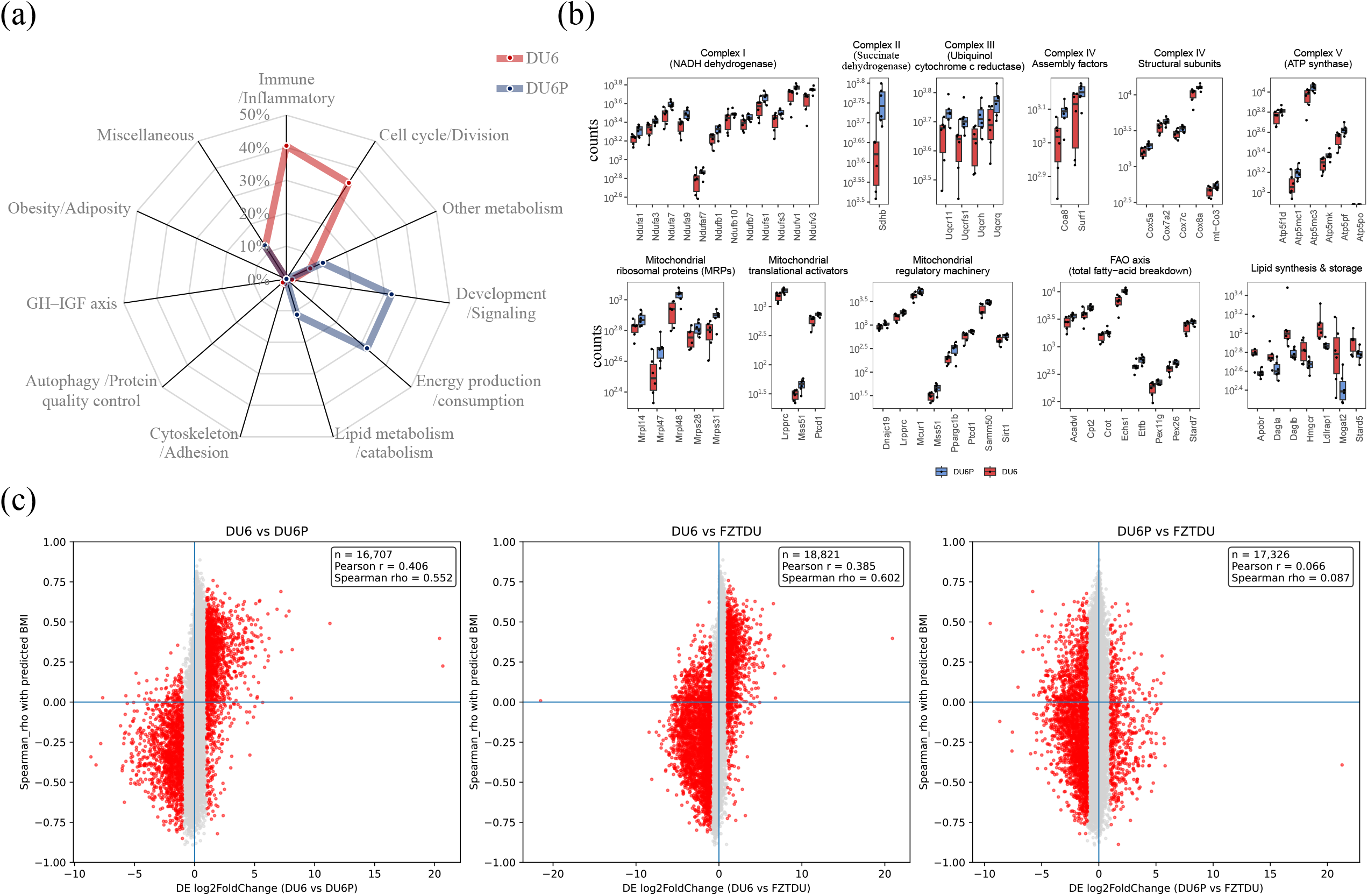
Comparative functional enrichment and mitochondrial pathway profiles in Titan mouse lines. (a) Radar plot showing the relative enrichment contribution of functional categories between DU6 and DU6P lines. Categories include Immune/Inflammatory, cell cycle/division, other metabolism, development/signaling, energy production/consumption, lipid metabolism/catabolism, cytoskeleton/adhesion, autophagy/protein quality control, GH–IGF axis, obesity/adiposity, and miscellaneous. DU6 and DU6P exhibit distinct enrichment patterns, highlighting differential metabolic and signaling profiles. The relative enrichment contribution of each category was calculated from the proportional sum of NES values of functions within that category relative to the total NES. (b) Boxplots of representative mitochondrial pathways and related processes. Shown are enrichment counts for oxidative phosphorylation complexes (I–V), mitochondrial translation, glucose and amino acid catabolism, lipid synthesis and storage, and organelle quality control. Comparisons reveal a marked shift toward oxidative and catabolic pathways in DU6P, contrasting with the more anabolic and lipid-storage–oriented profile of DU6. (c) Scatter plots showing gene-level concordance between gene-expression correlation with predicted BMI and differential expression across pairwise comparisons among DU6, DU6P, and FZTDU. The x axis indicates differential expression (log2 fold change), and the y axis shows the corresponding Spearman correlation coefficient with predicted BMI. Genes with |log2 fold change| < 1 are shown in light grey, and genes with |log2 fold change| ≥ 1 are shown in red. The vertical line indicates log2 fold change of zero; and the horizontal line marks zero correlation.

Because GSEA captures distributed transcriptome-wide signals, we next performed differentially expressed gene (DEG) enrichment analysis to obtain sharper resolution of functional modules (Fig. 3b, Table S1 and Supplementary Method). Genes identified by DESeq2 with an adjusted p-value < 0.05 were defined as DEGs. This analysis revealed opposing metabolic programs between the two lines. In DU6, lipid synthesis and storage pathways were strongly enriched (OR *≈* 76.5, p = 3.8 × 10^−11^), consistent with an anabolic program favoring lipid accumulation. In DU6P, enrichment was dominated by oxidative metabolism, including multiple respiratory chain complexes: Complex I (OR *≈* 32.1, p = 1.2 × 10^−13^), Complex II (OR *≈* 27.1, p = 0.048), Complex III (OR *≈* 47.1, p = 6.7 × 10^−6^), Complex IV structural components (OR *≈* 31.8, p = 2.0 × 10^−6^) and assembly factors (OR *≈* 10.9, p = 0.018), and Complex V/ATP synthase (OR *≈* 27.7, p = 3.5 × 10^−7^). DU6P was also enriched in mitochondrial ribosomal proteins (OR *≈* 5.65, p = 0.0027), mitochondrial translational activators (OR *≈* 30.7, p = 2.8 × 10^−4^), and mitochondrial regulatory machinery (OR *≈* 15.3, p = 2.1 × 10^−7^), defining a strong OXPHOS-centered program. Furthermore, DU6P exhibited expanded fatty acid oxidation (FAO) capacity, encompassing mitochondrial FAO enzymes, ETF coupling factors, and peroxisomal biogenesis (OR *≈* 20.4, p = 2.9 × 10^−8^), underscoring a broad oxidative orientation.

Analysis of the growth hormone (GH) signaling axis provided an additional discriminant feature. While the strict GH core set showed no significant difference between lines, the extended GH axis— incorporating downstream metabolic effectors—was significantly enriched in DU6P (OR *≈* 13.7, p = 3.5 × 10^−4^) but not in DU6 (OR *≈* 11.2, p = 0.090). This suggests that GH-axis-associated transcriptional regulation is more strongly represented in DU6P and may be linked to its enhanced oxidative program.

To test whether these opposing transcriptional states were also aligned with the human-trained BMI axis at the gene level, we compared differential-expression effect sizes with gene-wise correlations to predicted BMI across all three pairwise mouse-line contrasts (Fig. 3c). Strong positive concordance was observed for contrasts involving DU6: log2 fold changes were positively associated with Spearman correlation coefficients in both DU6 vs DU6P (Pearson r = 0.406; Spearman rho = 0.552) and DU6 vs FZTDU (Pearson r = 0.385; Spearman rho = 0.602). In contrast, DU6P vs FZTDU showed only minimal association (Pearson r = 0.066; Spearman rho = 0.087). This pattern indicates that the predicted-BMI axis does not simply reflect generic inter-line divergence, but preferentially tracks the molecular program associated with the DU6 state. In this framework, DU6 corresponds to the obesity-vulnerable, lipid-anabolic pole of the model, whereas DU6P corresponds to an oxidative-catabolic and metabolically resilient counter-state.

Taken together, these analyses identify DU6 and DU6P as two opposing adipose transcriptional configurations: one biased toward lipid accumulation, inflammatory remodeling, and obesity vulnerability, and the other toward mitochondrial function, substrate oxidation, and metabolic resilience. The strong alignment of the predicted-BMI axis with contrasts involving DU6 further supports the conclusion that this cross-species transcriptomic model captures a biologically meaningful obesity-related state rather than reflecting nonspecific inter-line differences.

### Cross-species transcriptomics model uncovers protective mitochondrial programs distinguishing MHO from MUO

To explore the utility of our transcriptome-based model in understanding obesity heterogeneity, we examined the relationship between predicted and observed BMI across patients. Comparing molecularly derived predictions with phenotypic BMI enabled stratification into metabolically distinct subgroups (Fig. 4a). This approach identified individuals whose gene expression profiles diverged from their physical BMI burden. Patients assigned to the MHO group exhibited molecular signatures indicative of reduced metabolic risk despite elevated BMI, whereas those in the MUO group displayed expression patterns consistent with adverse metabolic remodeling.

**Figure 4.**
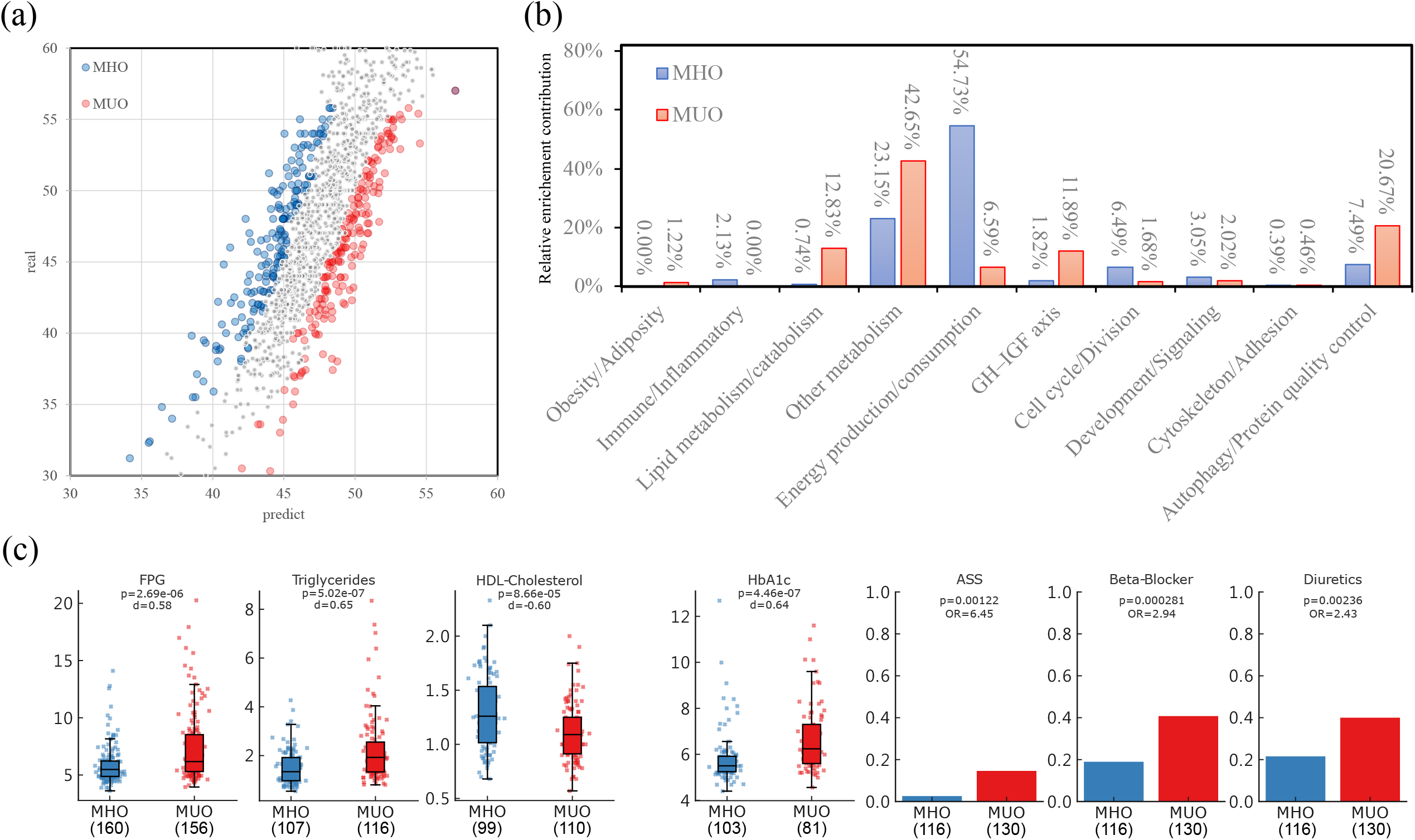
Molecular prediction of obesity subtypes and functional enrichment profiles. (a) Scatter plot comparing predicted versus observed BMI across individuals. Patients were stratified into MHO (blue) and MUO (red) groups based on a post-hoc LOWESS model, illustrating clear divergence between molecular and phenotypic obesity states. (b) Bar chart summarizing the relative enrichment contribution of functional categories for MHO and MUO subgroups. MHO is predominantly associated with energy production/consumption, whereas MUO shows higher contributions from other metabolism and autophagy/protein quality control pathways, highlighting distinct molecular mechanisms underlying obesity heterogeneity. The relative enrichment contribution of each functional category was calculated with the miscellaneous category excluded. (c) After exclusion of three outliers (vis-2520, the MUO individual with the highest gGT; vis-2504, the MHO individual with the highest gGT; and vis-2149, the MUO individual with the highest Triglycerides), seventy-seven clinical and biochemical parameters were compared between MHO (blue) and MUO (red) groups. Binary variables were analyzed using Fisher’s exact test, and continuous variables were assessed using the Mann– Whitney U test. Effect sizes are reported as odds ratios (ORs) for binary variables—indicating the relative likelihood of a clinical feature in MUO compared with MHO—and as Cohen’s d for continuous variables, quantifying standardized differences in group means. Significant differences (FDR-adjusted q < 0.05 and Cohen’s d > 0.4) revealed higher FPG, triglycerides, and HbA1c, along with more frequent use of ASS, beta blockers, and diuretics in MUO, whereas HDL-cholesterol was elevated in MHO. These physiological and biochemical distinctions align with the transcriptome-based stratification, supporting metabolic divergence between MHO and MUO.

Analysis of MHO-upregulated genes revealed enrichment dominated by Energy production/consumption (55%), suggesting strong signals in mitochondrial energetics, including oxidative phosphorylation and ATP synthesis–coupled electron transport (Fig. 4b and Fig. S5). Secondary contributions were categorized as other metabolism, including RNA biosynthesis, intracellular sodium ion homeostasis, and nucleic acid metabolism. Together, these results suggest that MHO patients maintain an enhanced capacity for mitochondrial energy production and metabolism, reflecting coordinated growth and metabolic efficiency.

In contrast, the MUO profile displayed a distinct enrichment pattern characterized by broad transcriptional remodeling (Fig. 4b and Fig. S6). The strongest enrichment was observed in Other metabolism, which included RNA processing and oxoacid and nucleotide metabolism. Substantial signals were also detected in Autophagy/Protein quality control, reflecting activation of proteostatic and stress-adaptive mechanisms, and in Lipid metabolism/catabolism, represented by pathways such as fatty acid β-oxidation. Collectively, these results indicate that MUO-upregulated genes primarily participate in metabolic regulation, proteostasis, and lipid catabolism that support stress adaptation and tissue remodeling.

Moreover, clinical parameters were broadly consistent with these molecular distinctions (Fig. 4c). MHO individuals exhibited significantly lower fasting plasma glucose (FPG, p = 2.7 × 10^−6^) and triglycerides (p = 5.0 × 10^−7^), along with higher HDL-cholesterol (p = 8.7 × 10^−5^), indicating their more favorable glycemic control and lipid handling. In contrast, MUO participants showed higher glycated hemoglobin (HbA1c, p = 4.5 × 10^−7^). Cardiometabolic medication use was also more frequent in MUO, including acetylsalicylic acid (ASS, OR = 6.45, p = 0.0012), beta blockers (OR = 2.94, p = 0.00028), and diuretics (OR = 2.43, p = 0.0024). Together, these findings support the transcriptome-based stratification and further highlight the physiological and metabolic heterogeneity within obesity.

Together, these findings highlight a fundamental transcriptional divergence between MHO and MUO. MHO is characterized by mitochondrial energetics and translational upregulation, supporting a more efficient and regulated metabolic state, whereas MUO is marked by protein modification, lipid catabolism, and growth factor signaling, coupled with stress-responsive autophagy, indicative of a dysregulated and metabolically vulnerable phenotype.

Complementary to the transcriptome-based comparison between the human and mouse obesity states, we calculated the ROH to examine the relevance of molecular signatures to signatures of long-term selection. Genes whose scores belong to the top 10% based on their normalized scores were classified as high-ROH loci and were identified as regions with extensive allelic fixation and potential selection pressure. In the DU6 genome, 2,816 high-ROH genes were identified proving its consistency with the line’s long-term selection. Notably, only 41 genes were found to overlap with the differentially expressed genes (DU6 vs FZTDU; odds ratio = 0.34, p = 2.52 × 10^−14^) from the adipose tissue transcriptome reflecting the convergence of genetic fixation and transcriptional activation for some metabolic and growth-related pathways (Fig 5).

**Figure 5.**
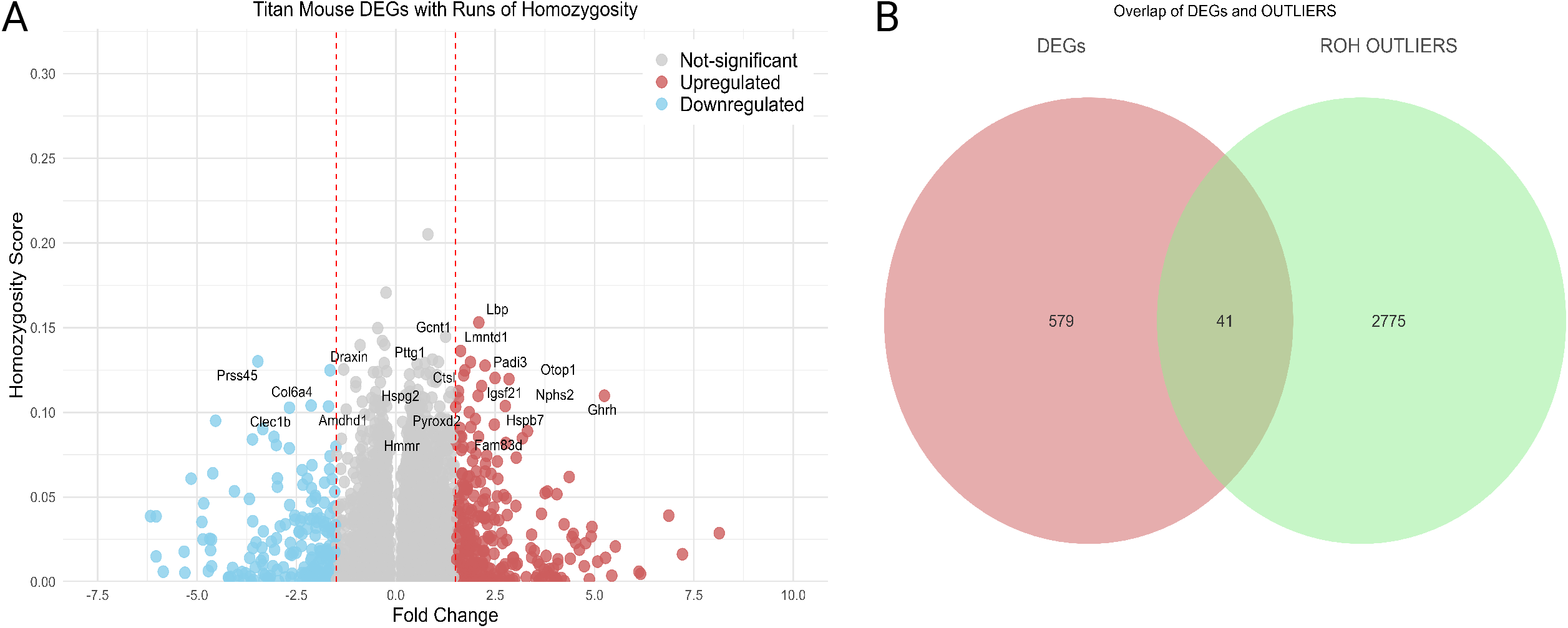
Partial convergence of transcriptional regulation and genomic fixation in Titan mice. (a) The volcano plot shows the fold change and homozygosity scores of the genes in Titan mice. The points represent genes and their direction regulation: upregulated (red) and downregulated (blue) while only the DEGs with elevated ROH scores are annotated and highlighted. Only few observable subsets of genes demonstrated both dysregulation and elevation in ROH implicating them as potential points of functional convergence between genomic fixation and phenotypic adaptation. (b) The diagram shows the overlap (n = 41) between the DEGs and ROH outlier genes. Altogether, these results imply that a portion of the ROH regions capturing selection signature relates and converges with detectable transcriptional changes highlighting a focused set of candidate genes contributing to the Titan mice phenotype.

One of the overlapping genes is the growth hormone-releasing hormone (*Ghrh*). The accumulation of homozygosity around this gene further supports the idea of allelic variant fixation linking to GH axis consistent with previously observed signaling differences.

Consistent with this dichotomy, the opposing metabolic configurations observed in the DU6P and DU6 mouse lines and the long-term genetic fixation observed in the DU6 genome point to a conserved molecular axis linking mitochondrial efficiency to metabolic resilience and anabolic remodeling to vulnerability. This cross-species coherence reinforces the generalizable nature of the transcriptome-based model and its capacity to capture evolutionarily conserved principles of metabolic health.

## Discussion

In this study we set out to determine whether conserved transcriptomic programs differentiate metabolically unhealthy from healthier obesity states. Our study establishes a transcriptome-based framework for dissecting the molecular underpinnings of obesity across human and mouse models. By integrating adipose tissue gene expression with machine learning, we demonstrated that transcriptomic features capture BMI-linked biological variation while preserving meaningful rank structure and revealing functional programs central to obesity biology. Our results are consistent with human adipose transcriptomic work showing that metabolic health is a BMI-independent molecular dimension: bulk visceral adipose tissue profiling can stratify obese individuals into metabolically healthier vs unhealthier groups despite similar adiposity^44,45^. This separation may also reflect a continuum across stages of metabolic dysregulation^46^. In addition, compact transcriptomic signature models linking MUO to comorbidities (e.g., MASLD using liver/VAT expression) highlight the translational potential of expression-based risk subtyping^47^. Furthermore, our transcriptomic model captures signatures spanning immune activation, lipid and energy metabolism, and cellular growth regulation, highlighting the multifactorial transcriptional basis of adiposity. Importantly, cross-species validation confirmed that the model reflects adiposity-specific molecular signals rather than gross body size, since it could distinguish between DU6 and DU6P mouse lines that are similar in body weight to length ratio, but highly divergent in adipose tissue phenotype. Several other studies have used cross-species approaches to identify genes associated with obesity and fat deposition^48-50^. Liu et al compared different machine learning methods to predict genes related to fat deposition in pigs, Acharjee et al. used publicly available human and mouse transcriptome data and machine learning to find obesity-associated genes that are common across species and Wang et al. integrated the skeletal muscle transcriptome of obese pigs with publicly available transcriptional profiles and ATAC-seq human and mouse data.

A major strength of our approach lies in its ability to bridge anthropometric measurements with molecular signatures. Traditional metrics such as BMI or body weight provide only crude estimates of adiposity and fail to distinguish between metabolically healthy and unhealthy states. Our transcriptome-based model overcomes this limitation by identifying protective versus pathogenic molecular programs, including enhanced mitochondrial energetics in MHO and remodeling/autophagy in MUO. This capacity to extract biologically meaningful differences from transcriptomic data offers potential for improved stratification of obesity subtypes and for guiding precision interventions.

Nevertheless, certain limitations must be acknowledged. First, BMI was chosen as the predictive label due to its availability and clinical relevance, but it is an imperfect surrogate for adiposity and metabolic health^8,25,51^. Future iterations of the model would benefit from incorporating additional phenotypes such as insulin sensitivity, lipid profiles, or imaging-derived fat distribution. Second, our analysis focused primarily on adipose tissue transcriptomes, while obesity is a systemic condition involving multiple tissues and inter-organ communication^52-54^. Expanding to cross-tissue or single-cell data would further refine mechanistic insights. Finally, although cross-species validation strengthens the robustness of the model, differences in human and mouse physiology necessitate careful translation of findings^15,55,56^.

DU6 mice are a unique non-inbred mouse model for polygenic obesity^17,57^. While the cross-species transcriptome model already demonstrated that the DU6 mouse line exhibits molecular features for MUO, the analysis for runs of homozygosity (ROH) further revealed information of its long-term genetic consolidation and suggests that persistent selection for growth traits can affect metabolic vulnerability at the genomic level. The fixation found in *Ghrh* alleles represents a compelling instance of this finding. Being both differentially expressed and having high homozygosity score, we can deduce that selection for body mass in DU6 has favored variants that enhance growth signalling, resulting in transcriptomic changes that ultimately promote lipid deposition and diminished oxidative capacity.

Looking forward, several avenues emerge. Functionally, the mitochondrial and protein translational pathways identified in MHO represent attractive targets for therapeutic intervention or biomarker development. This is supported by transcriptomic studies identifying coordinated OXPHOS gene-set downregulation in human metabolic disease^58^ and showing that short-term high-fat feeding suppresses mitochondrial/OXPHOS genes as an early molecular signature of diet-induced metabolic stress^59^. In parallel, translational control has emerged as a modifiable layer of adipose remodeling: PPARγ agonism can remodel translational machinery in adipose progenitors, but this response is impaired in obesity^60^. Methodologically, integrating transcriptomic predictors with other omics layers—such as epigenomics, proteomics, or metabolomics—could enhance predictive power and mechanistic resolution. Clinically, applying this framework to longitudinal cohorts may allow early identification of patients at risk for progression from metabolically healthy to unhealthy obesity, consistent with long-term prospective evidence that many initially metabolically healthy individuals transition to an unhealthy phenotype and that this conversion is accompanied by higher incident cardiovascular risk^61^. This view is reinforced by cohort analyses treating metabolic status as time-varying—highlighting the dynamic nature of “metabolic health” within obesity over extended follow-up^62^—and by detailed trajectory work describing transitions from healthy to unhealthy obesity and identifying clinical factors associated with conversion^63^. A systematic review/meta-analysis further synthesizes that MHO*→M*UO transition is common and that transitioners exhibit elevated CVD incidence, underscoring the value of early risk stratification and prevention windows^64^.

In a broader perspective, this study underscores the utility of cross-species transcriptomics for disentangling the complex biology of obesity. By aligning human and mouse molecular states, we reveal conserved programs that distinguish protective from deleterious adiposity. Beyond obesity, this strategy may be generalized to other complex diseases where phenotypic heterogeneity complicates diagnosis and treatment. Ultimately, transcriptome-informed classification holds promise for moving obesity research and clinical care toward a more mechanistic and personalized paradigm.

## Conclusions

We established a BMI-anchored transcriptomic framework that captures adiposity-linked molecular programs beyond gross anthropometric variation. Cross-species application showed that this framework reflects adiposity-specific transcriptional states rather than body size alone, distinguishing lipid-anabolic from oxidative-catabolic obesity phenotypes in mouse models. In humans, it resolved metabolically distinct obese subgroups, linking MHO to mitochondrial and translational programs associated with resilience, and MUO to stress-adaptive, catabolic, and remodeling pathways associated with vulnerability. These findings support a conserved molecular axis underlying obesity heterogeneity and provide a basis for more biologically informed classification of metabolic disease.

## Declarations

### Ethics approval and consent to participate

The LOBB study was approved by the Ethics Committee of the University of Leipzig (approval numbers: 159-12-21052012 and 017-12ek) and performed in accordance with the Declaration of Helsinki, the Bioethics Convention (Oviedo), and EU Directive on Clinical Trials (Directive 2001/20/EC). Written informed consent was obtained from all donors. All donors have been informed of the purpose, risks and benefits of the biobank. Ethical guidelines and EU legislation for privacy and confidentiality in personal data collection and processing is being followed, in particular directive 95/46/EC.

### Consent for publication

Not applicable.

### Availability of data and materials

Bulk RNAseq data from the LOBB reported in this study cannot be deposited in a public repository due to restrictions by patient consent. These restrictions are due to local data protection regulation in the written informed consent form tissue donors signed before taking part in the study. Access to human adipose tissue biobank data is regulated by the LOBB steering committee. Use of data is strictly limited to research purposes and not intended for commercialization. To request access, contact Matthias Blüher or Anne Hoffmann.

### Authors’ contributions

Conceptualization: CCL, YYS

Data curation: AH; AG

Formal analysis: AH, CCL, CYC, YYS

Funding acquisition: CCL, MB

Investigation: CCL, CYC, YYS

Methodology: CCL, CYC, YYS

Project administration: CCL

Resources: CCL, CW, MB

Supervision: CCL

Visualization: CCL, CYC, YYS

Writing – original draft: CCL, CYC, YYS

Writing – review & editing: AH, AG, CCL, CW, MB, YYS

## References

1 Tahir, U. A. & Gerszten, R. E. Molecular Biomarkers for Cardiometabolic Disease: Risk Assessment in Young Individuals. Circ Res 132, 1663–1673, doi:10.1161/CIRCRESAHA.123.322000 (2023).

2 World Health, O. The top 10 causes of death. (2024).

3 Collaborators, G. B. D. C. R. F. The global burden of cancer attributable to risk factors, 2010-19: a systematic analysis for the Global Burden of Disease Study 2019. Lancet 400, 563–591, doi:10.1016/S0140-6736(22)01438-6 (2022).

4 Livingston, G. et al. Dementia prevention, intervention, and care: 2024 report of the Lancet standing Commission. Lancet 404, 572–628, doi:10.1016/S0140-6736(24)01296-0 (2024).

5 Ndumele, C. E. et al. Cardiovascular-Kidney-Metabolic Health: A Presidential Advisory From the American Heart Association. Circulation 148, 1606–1635, doi:10.1161/CIR.0000000000001184 (2023).

6 Franks, P. W. & Sargent, J. L. Diabetes and obesity: leveraging heterogeneity for precision medicine. Eur Heart J 45, 5146–5155, doi:10.1093/eurheartj/ehae746 (2024).

7 Schulze, M. B. & Stefan, N. Metabolically healthy obesity: from epidemiology and mechanisms to clinical implications. Nat Rev Endocrinol 20, 633–646, doi:10.1038/s41574-024-01008-5 (2024).

8 World Health, O. Obesity and overweight. (2024).

9 Borgeson, E., Tavajoh, S., Lange, S. & Jessen, N. The challenges of assessing adiposity in a clinical setting. Nat Rev Endocrinol 20, 615–626, doi:10.1038/s41574-024-01012-9 (2024).

10 Rubino, F. et al. Definition and diagnostic criteria of clinical obesity. Lancet Diabetes Endocrinol 13, 221–262, doi:10.1016/S2213-8587(24)00316-4 (2025).

11 Lean, M. E. et al. Primary care-led weight management for remission of type 2 diabetes (DiRECT): an open-label, cluster-randomised trial. Lancet 391, 541–551, doi:10.1016/S0140-6736(17)33102-1 (2018).

12 Taylor, R. & Holman, R. R. Normal weight individuals who develop type 2 diabetes: the personal fat threshold. Clin Sci (Lond) 128, 405–410, doi:10.1042/CS20140553 (2015).

13 Taylor, R. et al. Aetiology of Type 2 diabetes in people with a ‘normal’ body mass index: testing the personal fat threshold hypothesis. Clin Sci (Lond) 137, 1333–1346, doi:10.1042/CS20230586 (2023).

14 Lean, M. E. et al. 5-year follow-up of the randomised Diabetes Remission Clinical Trial (DiRECT) of continued support for weight loss maintenance in the UK: an extension study. Lancet Diabetes Endocrinol 12, 233–246, doi:10.1016/S2213-8587(23)00385-6 (2024).

15 Borgeson, E., Boucher, J. & Hagberg, C. E. Of mice and men: Pinpointing species differences in adipose tissue biology. Front Cell Dev Biol 10, 1003118, doi:10.3389/fcell.2022.1003118 (2022).

16 Loft, A. et al. Towards a consensus atlas of human and mouse adipose tissue at single-cell resolution. Nat Metab 7, 875–894, doi:10.1038/s42255-025-01296-9 (2025).

17 Muller-Eigner, A. et al. Dietary intervention improves health metrics and life expectancy of the genetically obese Titan mouse. Commun Biol 5, 408, doi:10.1038/s42003-022-03339-3 (2022).

18 Gille, B. et al. Titan mice as a model to test interventions that attenuate frailty and increase longevity. Geroscience 46, 3599–3606, doi:10.1007/s11357-023-01045-4 (2024).

19 Gjermeni, E. et al. The impact of dietary interventions on cardiometabolic health. Cardiovasc Diabetol 24, 234, doi:10.1186/s12933-025-02766-w (2025).

20 Sawitzky, M. et al. Phenotype selection reveals coevolution of muscle glycogen and protein and PTEN as a gate keeper for the accretion of muscle mass in adult female mice. PLoS One 7, e39711, doi:10.1371/journal.pone.0039711 (2012).

21 Gjermeni, E. et al. Obesity-An Update on the Basic Pathophysiology and Review of Recent Therapeutic Advances. Biomolecules 11, doi:10.3390/biom11101426 (2021).

22 Khawaja, T., Nied, M., Wilgor, A. & Neeland, I. J. Impact of Visceral and Hepatic Fat on Cardiometabolic Health. Curr Cardiol Rep 26, 1297–1307, doi:10.1007/s11886-024-02127-1 (2024).

23 Kubota, S. & Yabe, D. Visceral Adipose Tissue Quality and its Impact on Metabolic Health. J Clin Endocrinol Metab 109, e1665–e1666, doi:10.1210/clinem/dgae021 (2024).

24 Lee, M. J. & Kim, J. The pathophysiology of visceral adipose tissues in cardiometabolic diseases. Biochem Pharmacol 222, 116116, doi:10.1016/j.bcp.2024.116116 (2024).

25 Bluher, M. Metabolically Healthy Obesity. Endocr Rev 41, doi:10.1210/endrev/bnaa004 (2020).

26 Kloting, N. et al. Insulin-sensitive obesity. Am J Physiol Endocrinol Metab 299, E506–515, doi:10.1152/ajpendo.00586.2009 (2010).

27 Langhardt, J. et al. Effects of Weight Loss on Glutathione Peroxidase 3 Serum Concentrations and Adipose Tissue Expression in Human Obesity. Obes Facts 11, 475–490, doi:10.1159/000494295 (2018).

28 Picelli, S. et al. Full-length RNA-seq from single cells using Smart-seq2. Nat Protoc 9, 171–181, doi:10.1038/nprot.2014.006 (2014).

29 Chen, S., Zhou, Y., Chen, Y. & Gu, J. fastp: an ultra-fast all-in-one FASTQ preprocessor. Bioinformatics 34, i884–i890, doi:10.1093/bioinformatics/bty560 (2018).

30 Dobin, A. et al. STAR: ultrafast universal RNA-seq aligner. Bioinformatics 29, 15–21, doi:10.1093/bioinformatics/bts635 (2013).

31 Liao, Y., Smyth, G. K. & Shi, W. featureCounts: an efficient general purpose program for assigning sequence reads to genomic features. Bioinformatics 30, 923–930, doi:10.1093/bioinformatics/btt656 (2014).

32 Renne, U. et al. Lifelong obesity in a polygenic mouse model prevents age- and diet-induced glucose intolerance-obesity is no road to late-onset diabetes in mice. PLoS One 8, e79788, doi:10.1371/journal.pone.0079788 (2013).

33 Schueler, L. Mouse strain Fzt:DU and its use as model in animal breeding research. Archiv für Tierzucht 28, 357–363 (1985).

34 Dietl, G., Langhammer, M. & Renne, U. Model simulations for genetic random drift in the outbred strain Fzt: DU. Archiv für Tierzucht 47, 595–604 (2004).

35 Rankin, A. L. et al. IL-33 induces IL-13-dependent cutaneous fibrosis. J Immunol 184, 1526–1535, doi:10.4049/jimmunol.0903306 (2010).

36 Abràmoff, M. D., Magalhães, P. J. & Ram, S. J. Image processing with ImageJ. Biophotonics International 11, 36–42 (2004).

37 Galarraga, M. et al. Adiposoft: automated software for the analysis of white adipose tissue cellularity in histological sections. J Lipid Res 53, 2791–2796, doi:10.1194/jlr.D023788 (2012).

38 Vilella, A. J. et al. EnsemblCompara GeneTrees: Complete, duplication-aware phylogenetic trees in vertebrates. Genome Res 19, 327–335, doi:10.1101/gr.073585.107 (2009).

39 Harrison, P. W. et al. Ensembl 2024. Nucleic Acids Res 52, D891–D899, doi:10.1093/nar/gkad1049 (2024).

40 Subramanian, A. et al. Gene set enrichment analysis: a knowledge-based approach for interpreting genome-wide expression profiles. Proc Natl Acad Sci U S A 102, 15545–15550, doi:10.1073/pnas.0506580102 (2005).

41 Love, M. I., Huber, W. & Anders, S. Moderated estimation of fold change and dispersion for RNA-seq data with DESeq2. Genome Biol 15, 550, doi:10.1186/s13059-014-0550-8 (2014).

42 Shafer, A. B. A. & Kardos, M. Runs of Homozygosity and Inferences in Wild Populations. Mol Ecol 34, e17641, doi:10.1111/mec.17641 (2025).

43 Hewett, A. M., Stoffel, M. A., Peters, L., Johnston, S. E. & Pemberton, J. M. Selection, recombination and population history effects on runs of homozygosity (ROH) in wild red deer (Cervus elaphus). Heredity (Edinb) 130, 242–250, doi:10.1038/s41437-023-00602-z (2023).

44 Das, S. K., Ma, L. & Sharma, N. K. Adipose tissue gene expression and metabolic health of obese adults. Int J Obes (Lond) 39, 869–873, doi:10.1038/ijo.2014.210 (2015).

45 Zhou, Q. et al. Metabolic Health Status Contributes to Transcriptome Alternation in Human Visceral Adipose Tissue During Obesity. Obesity (Silver Spring) 28, 2153–2162, doi:10.1002/oby.22950 (2020).

46 Rosendo-Silva, D. et al. Clinical and molecular profiling of human visceral adipose tissue reveals impairment of vascular architecture and remodeling as an early hallmark of dysfunction. Metabolism 153, 155788, doi:10.1016/j.metabol.2024.155788 (2024).

47 Meroni, M. et al. A novel gene signature to diagnose MASLD in metabolically unhealthy obese individuals. Biochem Pharmacol 218, 115925, doi:10.1016/j.bcp.2023.115925 (2023).

48 Acharjee, A., Wijesinghe, S. N., Russ, D., Gkoutos, G. & Jones, S. W. Cross-species transcriptomics identifies obesity associated genes between human and mouse studies. J Transl Med 22, 592, doi:10.1186/s12967-024-05414-1 (2024).

49 Liu, H. et al. Using Machine Learning to Identify Biomarkers Affecting Fat Deposition in Pigs by Integrating Multisource Transcriptome Information. J Agric Food Chem 70, 10359–10370, doi:10.1021/acs.jafc.2c03339 (2022).

50 Wang, Y. et al. Obesity induced transcriptional changes in skeletal muscle across different species. PLoS One 20, e0327988, doi:10.1371/journal.pone.0327988 (2025).

51 Romero-Corral, A. et al. Accuracy of body mass index in diagnosing obesity in the adult general population. Int J Obes (Lond) 32, 959–966, doi:10.1038/ijo.2008.11 (2008).

52 Jia, Z. et al. Crosstalk between fat tissue and muscle, brain, liver, and heart in obesity: cellular and molecular perspectives. Eur J Med Res 29, 637, doi:10.1186/s40001-024-02176-w (2024).

53 Priest, C. & Tontonoz, P. Inter-organ cross-talk in metabolic syndrome. Nat Metab 1, 1177–1188, doi:10.1038/s42255-019-0145-5 (2019).

54 Wang, S. et al. Effects of multi-organ crosstalk on the physiology and pathology of adipose tissue. Front Endocrinol (Lausanne) 14, 1198984, doi:10.3389/fendo.2023.1198984 (2023).

55 Lempesis, I. G., Tsilingiris, D., Liu, J. & Dalamaga, M. Of mice and men: Considerations on adipose tissue physiology in animal models of obesity and human studies. Metabol Open 15, 100208, doi:10.1016/j.metop.2022.100208 (2022).

56 Delhanty, P. J. D. & Visser, J. A. Navigating the Strengths and Constraints of Mouse Models in Obesity Research. Endocrinology 166, doi:10.1210/endocr/bqaf123 (2025).

57 Palma-Vera, S. E. et al. Genomic characterization of the world’s longest selection experiment in mouse reveals the complexity of polygenic traits. BMC Biol 20, 52, doi:10.1186/s12915-022-01248-9 (2022).

58 Mootha, V. K. et al. PGC-1alpha-responsive genes involved in oxidative phosphorylation are coordinately downregulated in human diabetes. Nat Genet 34, 267–273, doi:10.1038/ng1180 (2003).

59 Sparks, L. M. et al. A high-fat diet coordinately downregulates genes required for mitochondrial oxidative phosphorylation in skeletal muscle. Diabetes 54, 1926–1933, doi:10.2337/diabetes.54.7.1926 (2005).

60 De Siqueira, M. K. et al. PPARgamma-dependent remodeling of translational machinery in adipose progenitors is impaired in obesity. Cell Rep 43, 114945, doi:10.1016/j.celrep.2024.114945 (2024).

61 Eckel, N. et al. Transition from metabolic healthy to unhealthy phenotypes and association with cardiovascular disease risk across BMI categories in 90 257 women (the Nurses’ Health Study): 30 year follow-up from a prospective cohort study. Lancet Diabetes Endocrinol 6, 714–724, doi:10.1016/S2213-8587(18)30137-2 (2018).

62 Hinnouho, G. M. et al. Metabolically healthy obesity and the risk of cardiovascular disease and type 2 diabetes: the Whitehall II cohort study. Eur Heart J 36, 551–559, doi:10.1093/eurheartj/ehu123 (2015).

63 Mendes, F. D. et al. From healthy to unhealthy obesity: A longitudinal study of adults in ELSA-Brasil. PLOS Glob Public Health 5, e0004325, doi:10.1371/journal.pgph.0004325 (2025).

64 Abiri, B., Koohi, F., Ebadinejad, A., Valizadeh, M. & Hosseinpanah, F. Transition from metabolically healthy to unhealthy overweight/obesity and risk of cardiovascular disease incidence: A systematic review and meta-analysis. Nutr Metab Cardiovasc Dis 32, 2041–2051, doi:10.1016/j.numecd.2022.06.010 (2022).

