## Supplementary Information for "A conserved transcriptomic model defines metabolic resilience and vulnerability in obesity"

### SI Materials and Methods

#### Functional mapping of enriched GO terms

To determine whether model-selected genes converged on biological processes relevant to obesity, we systematically mapped enriched GO terms from the GSEA analysis to four predefined functional themes: (1) Obesity/Adiposity, (2) Lipid metabolism/catabolism, (3) Energy production/consumption, and (4) GH–IGF axis regulation.

First, GSEA results derived from the model-selected genes were curated by retrieving all significantly enriched GO terms. Each term was then examined for biological relevance through keyword matching and mechanistic interpretation. Specifically, terms containing or associated with glucose, carbohydrate, adipose extracellular matrix remodeling, or inflammatory cytokines were assigned to the Obesity/Adiposity category. Terms directly involving lipid, fatty acid, acyl-CoA, phospholipid, glycolipid, or sphingolipid metabolism or transport were assigned to the Lipid metabolism/catabolism category. Processes related to mitochondria, pyruvate metabolism, acetyl-CoA production, GTP/ATP utilization, or oxidative phosphorylation were classified under Energy production/consumption. Finally, GO terms linked to MAPK/JNK cascades, steroid hormone receptor signaling, ERBB family pathways, or other endocrine growth factor responses were considered representative of the GH–IGF axis.

Where direct keyword matches were absent, functional associations were determined through established biological knowledge, focusing on known mechanisms of adipose tissue biology, systemic metabolism, and hormone regulation.

To improve interpretability beyond these four targets, terms not captured by the primary categories were further refined into broader biological systems. These included Immune/Inflammatory (e.g., interleukin regulation, antigen presentation, Toll-like receptor signaling), Cell cycle/Division (e.g., DNA replication, spindle assembly, checkpoint signaling), Autophagy/Protein quality control (e.g., autophagosome organization, ubiquitination, unfolded protein response), Cytoskeleton/Adhesion (e.g., actin and microtubule dynamics, extracellular matrix remodeling, junction organization), Development/Signaling (e.g., angiogenesis, Notch signaling, neuronal differentiation), and Other metabolism (e.g., nucleotide and polyamine metabolism). GO terms that did not clearly fall into any of these themes were assigned to a Miscellaneous category.

#### Functional categorization and enrichment of DEGs

To systematically evaluate the biological processes represented in the differentially expressed genes (DEGs) between DU6 and DU6P, we curated functional gene categories relevant to mitochondrial metabolism, lipid biology, and hormone signaling. Category definitions were assembled from a combination of established pathway databases (KEGG, Reactome, Gene Ontology, and MSigDB Hallmark collections) and literature-based expert curation. Gene symbols were harmonized to the official Mus musculus (mouse) nomenclature using the MGI database to ensure consistency. The definitions of each category are listed below and detailed gene list was recorded in the Supplementary Table S1.

**Respiratory chain complexes (Complexes I–V).** Subunit genes of the five oxidative phosphorylation complexes were obtained from the MitoCarta3.0 database and cross-referenced with KEGG OXPHOS pathway annotations. Complex I (NADH dehydrogenase) contained 44 nuclear-encoded subunits, Complex II (succinate dehydrogenase) 4 subunits, Complex III (ubiquinol–cytochrome c reductase) 11 subunits, Complex IV (cytochrome c oxidase) 18 structural subunits and 17 assembly factors, and Complex V (ATP synthase) 24 subunits.

**Mitochondrial translation machinery.** Nuclear-encoded mitochondrial ribosomal proteins (MRPs; 78 genes) and translational activators (11 genes including *Mss51, Lrpprc, Ptcd1*) were obtained from MitoCarta3.0 and recent mitochondrial translation studies.

**Mitochondrial regulatory machinery.** Genes regulating organelle biogenesis, calcium homeostasis, protein import, and transcriptional control (52 genes) were defined by merging mitochondrial biogenesis regulators (*Ppargc1b, Sirt1*), import machinery (TIM/SAM components such as *Dnajc19, Samm50*), calcium uniporter regulators (*Mcur1*), and mitochondrial translational activators.

**Fatty-acid oxidation (FAO) axis.** Genes encoding β-oxidation enzymes, electron transfer flavoproteins

(ETF), and peroxisomal fatty-acid metabolism proteins (41 genes) were collated from KEGG FAO pathway, MitoCarta3.0, and the PEX gene family. This unified "FAO axis" captured total fatty-acid breakdown capacity, including mitochondrial FAO (*Acadvl, Cpt2, Echs1*), ETF coupling (*Etfb*), and peroxisomal lipid metabolism (*Pex11g, Pex26*).

**Lipid synthesis and storage.** Genes involved in de novo fatty-acid synthesis, triglyceride biosynthesis, lipid droplet formation, and sterol/cholesterol handling (38 genes) were defined using Reactome lipid metabolism pathways and prior lipogenesis-focused studies. Examples include cholesterol synthesis (*Hmgcr*), mono-/diacylglycerol acyltransferases (*Mogat2, Dagla, Daglb*), lipoprotein handling (*Apobr, Ldlrap1*), and sterol transfer proteins (*Stard5*).

**Growth hormone (GH) signaling axis.** A "core" GH axis gene set (18 genes) was defined based on canonical GH–JAK–STAT signaling, including receptors (*Gh, Ghr*), downstream effectors (*Stat5a/b, Jak2, Socs2/3, Cish*), and insulin-like growth factor genes (*Igf1, Igfals, Igfbp1–6*). An "extended" GH axis (28 genes) was additionally defined to capture downstream metabolic mediators (*Igf2, Igf2r, Igf1r, Ppargc1a, Ppargc1b, Cebpa/b, Stat3, Socs1, Irs1*).

Finally, DEGs identified between DU6 and DU6P were intersected with each curated category using official gene symbols. For each category, enrichment was tested using a two-tailed Fisher's exact test, with the full set of 16,687 annotated mouse genes as background. For each comparison, we report the observed number of DEGs in the category, expected overlap by chance, odds ratio (OR), and *p*-value. This approach enabled quantification of overrepresentation for both broad mitochondrial pathways (OXPHOS, FAO) and focused functional modules (lipid synthesis, GH axis).

##### Independent histological characterization of adipose depots in DU6, DU6P, and FZTDU mice

To validate our findings at the transcriptional level, we performed histological analyses on epigonadal and inguinal WAT (Supplementary Fig. S2a,b). Adipocyte size measurement revealed that DU6 mice exhibited pronounced hypertrophy across both depots; epigonadal adipocytes were 2.3-fold larger than FZTDU controls ($p < 0.0001$) and 3.8-fold larger than those of DU6P ($p < 0.0001$). This hypertrophic phenotype was more evident in the inguinal depot, where DU6 cells were 4.9-fold larger than FZTDU ($p = 0.0010$) and 6.3-fold larger than DU6P ($p = 0.0007$). In contrast, DU6P mice maintained a lean cellular profile, with epigonadal adipocytes being significantly smaller than those of the FZTDU control group ($p = 0.0123$). Notably, in the inguinal depots of both FZTDU and DU6P mice, we identified clusters of multilocular cells resembling beige adipocytes (indicated by arrows in Supplementary Fig. S2b), which were absent in DU6 tissue, suggesting that high-fat selection may impair the browning potential of subcutaneous fat. Inflammatory infiltration was assessed via F4/80 immunohistochemical staining to visualize crown-like structures (CLS), which represent macrophages surrounding dying adipocytes (Supplementary Fig. S2c,d). In the epigonadal depot, no statistically significant differences in CLS number were observed between the three mouse lines. However, in the inguinal depot, DU6 mice displayed a significant increase in CLS compared to FZTDU, with the median number being 3.8-fold higher ($p = 0.0022$). This increase reflects a localized pro-inflammatory response associated with adipocyte size expansion seen in the DU6 line. Due to a lack of available tissue, only a single sample was analyzed for the DU6P inguinal group, preventing its inclusion in the formal statistical comparisons for this specific site.

**SI Figures**

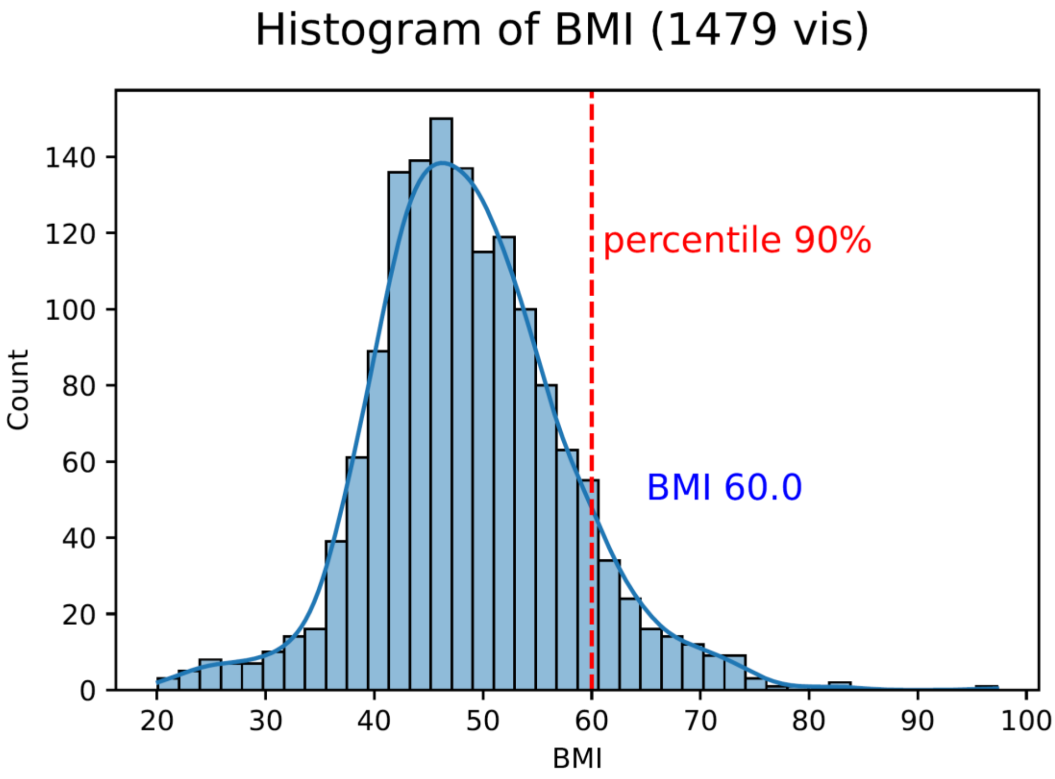

**Figure S1. BMI distribution in human visceral adipose tissue samples.**

Histogram of BMI values from 1,479 human visceral adipose tissue samples. The 90th percentile (red dashed line) corresponds to BMI ≈ 60.0. Sample counts by BMI range: <30 (n = 31), 30–40 (n = 159), 40–50 (n = 669), 50–60 (n = 470), ≥60 (n = 150).

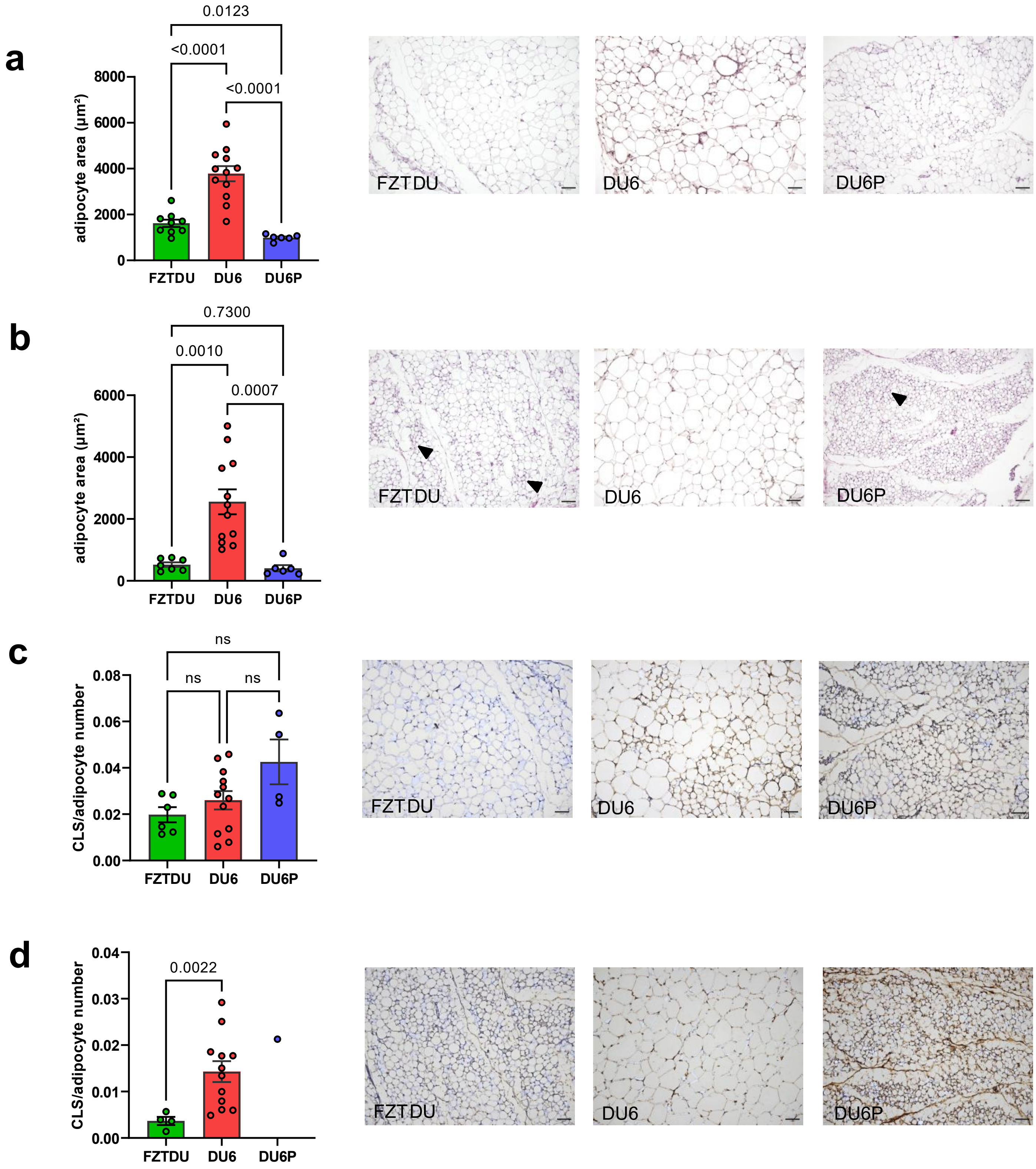

**Figure S2. Independent histological characterization of adipose depots in DU6, DU6P, and FZTDU mice.**

(a and b) Representative hematoxylin and eosin staining and adipocyte size quantification in epigonadal (a) and inguinal (b) white adipose tissue. (c and d) Representative F4/80 immunohistochemical staining and crown-like structure quantification in epigonadal (c) and inguinal (d) white adipose tissue. Analyses were performed in an independent cohort. ns, not significant.

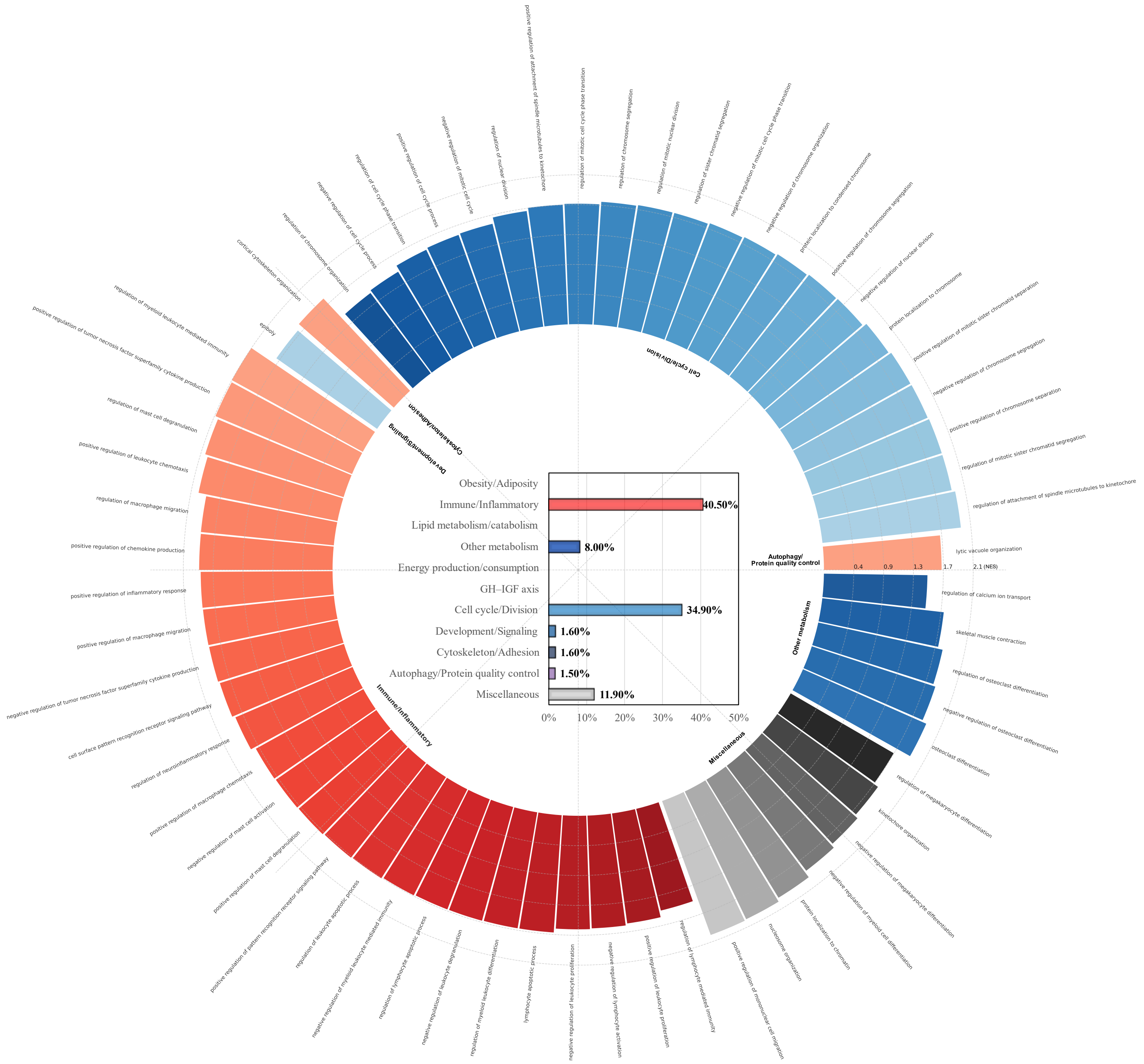

**Figure S3. Circular representation of relative enrichment contributions of individual functions grouped by biological category in DU6 mice.**

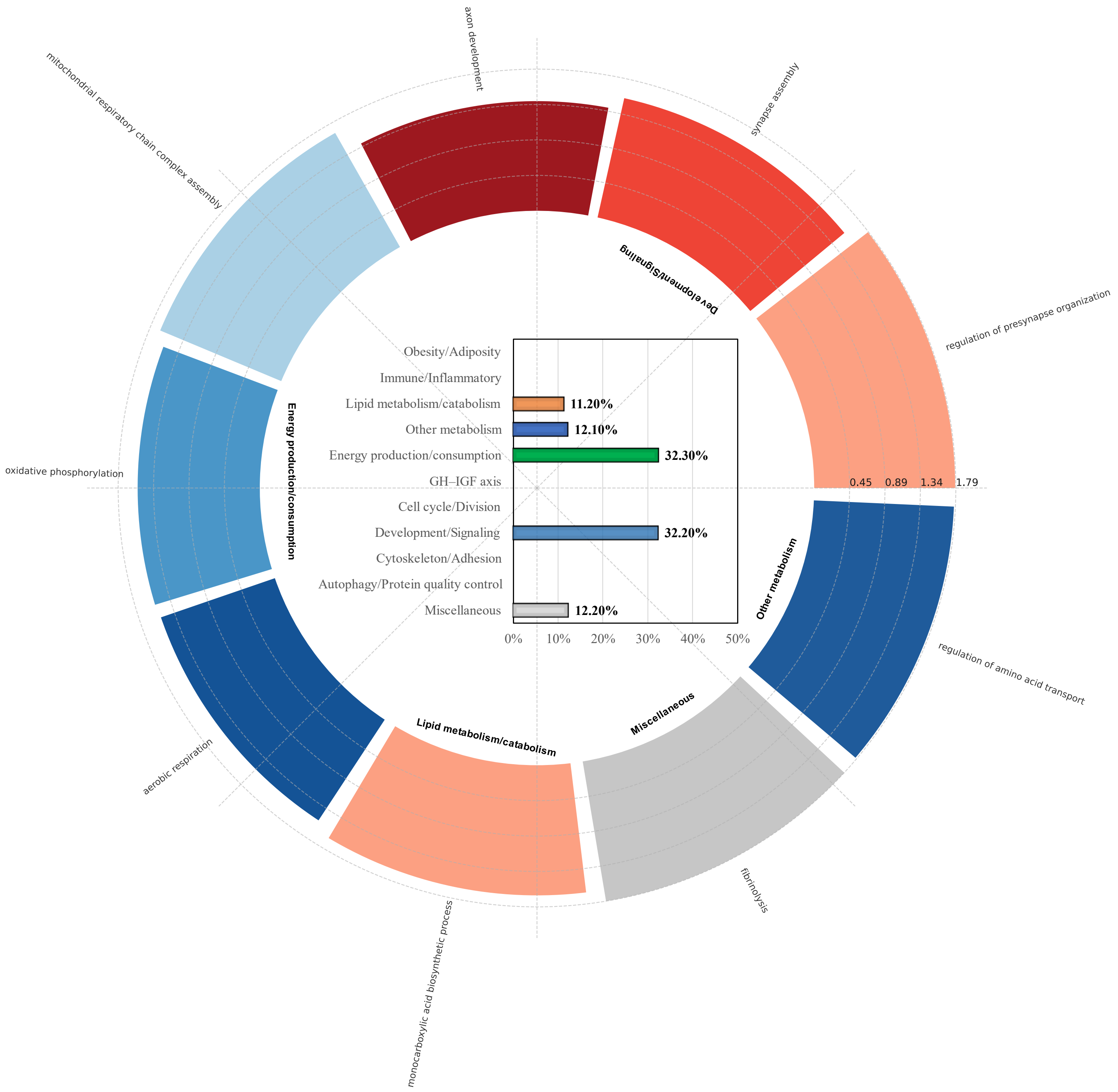

**Figure S4. Circular representation of relative enrichment contributions of individual functions grouped by biological category in DU6P mice.**

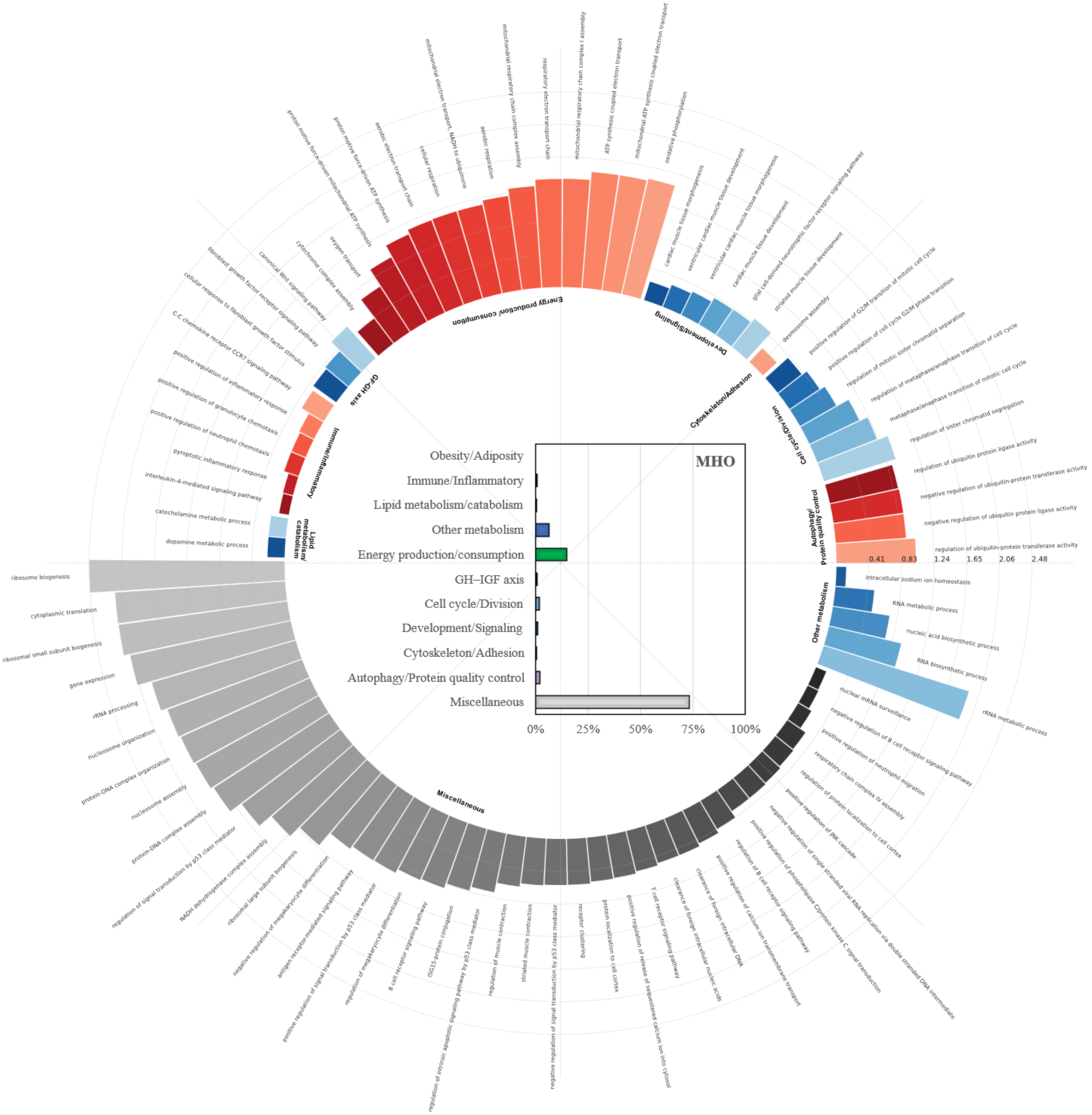

**Figure S5. Circular representation of relative enrichment contributions of individual functions grouped by biological category in MHO patients.**

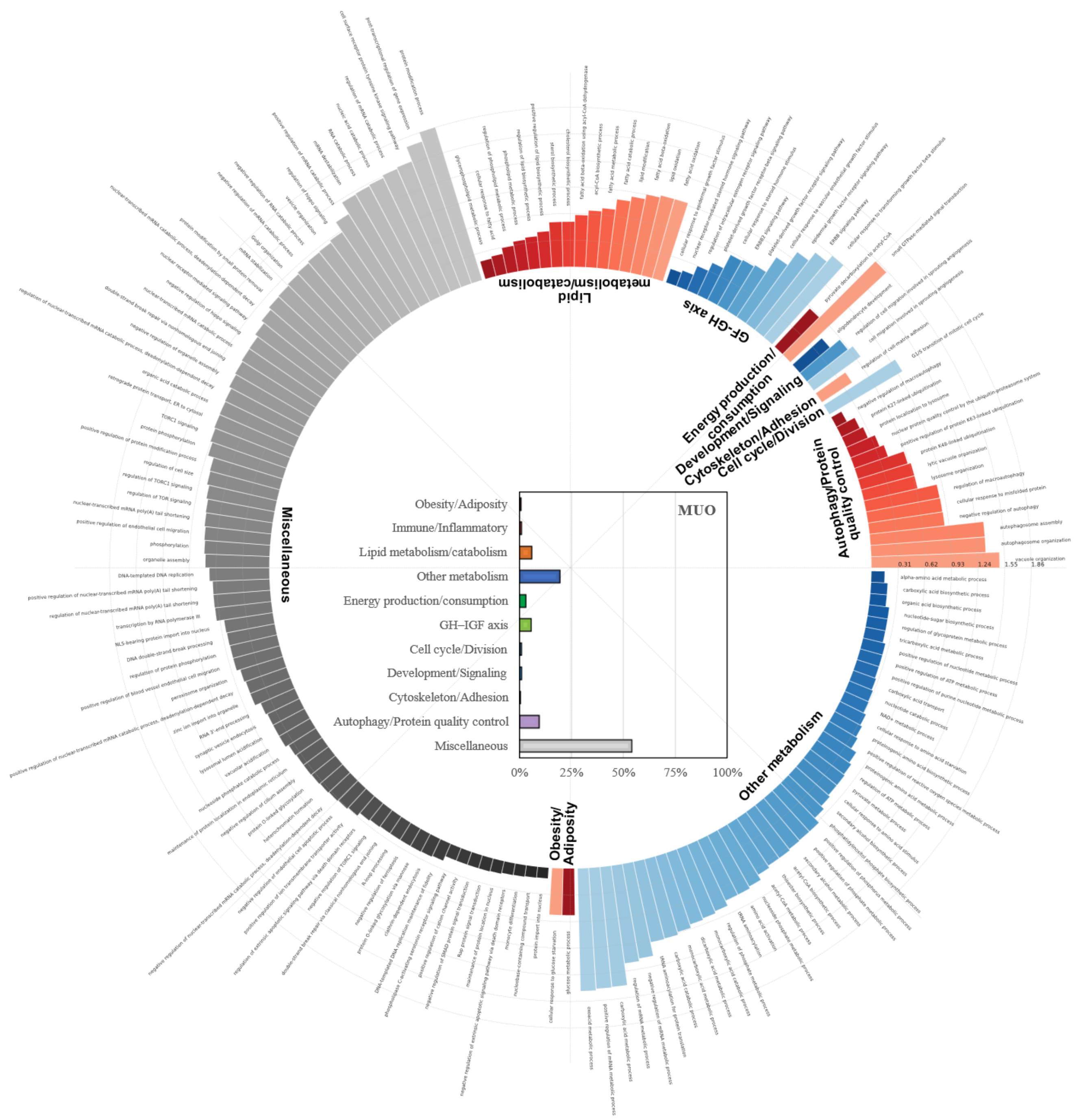

**Figure S6. Circular representation of relative enrichment contributions of individual functions grouped by biological category in MUO patients.**

#### SI Tables

**Table S1. Functional enrichment of mitochondrial complexes, regulatory machinery, and lipid metabolism pathways in DU6P and DU6 mouse lines.**

| Category | Odds Ratio | p-value | Observed Genes |
|---|---|---|---|
| | **DU6P** | | |
| Complex I (NADH dehydrogenase) | 32.1 | $1.2 \times 10^{-13}$ | *Ndufa1, Ndufa3, Ndufa7, Ndufa9, Ndufb1, Ndufb7, Ndufb10, Ndufs1, Ndufs3, Ndufv1, Ndufv3, Ndufaf7* |
| Complex II (Succinate dehydrogenase) | 27.1 | $4.8 \times 10^{-2}$ | *Sdhb* |
| Complex III (Ubiquinol–cytochrome c reductase) | 47.1 | $6.7 \times 10^{-6}$ | *Uqcrfs1, Uqcrq, Uqcr11, Uqcrh* |
| Complex IV – Assembly factors | 10.9 | $1.8\times 10^{-2}$ | *Surf1, Coa8* |
| Complex IV – Structural subunits | 31.8 | $2.0 \times 10^{-6}$ | *Cox5a, Cox7a2, Cox7c, Cox8a, mt-Co3* |
| Complex V (ATP synthase) | 27.7 | $3.5 \times 10^{-7}$ | *Atp5f1d, Atp5mc1, Atp5mc3, Atp5mk, Atp5pf, Atp5po* |
| Mitochondrial ribosomal proteins (MRPs) | 5.65 | $2.7\times 10^{-3}$ | *Mrpl14, Mrpl47, Mrpl48, Mrps28, Mrps31* |
| Mitochondrial translational activators | 30.7 | $2.8 \times 10^{-4}$ | *Lrpprc, Mss51, Ptcd1* |
| Mitochondrial regulatory machinery (merged: translational activators + biogenesis regulators + import machinery + MCU regulators) | 15.3 | $2.1 \times 10^{-7}$ | *Mss51, Lrpprc, Ptcd1, Ppargc1b, Sirt1, Dnajc19, Samm50, Mcur1* |
| FAO axis (total fatty-acid breakdown) (FAO + ETF + PEX) | 20.4 | $2.9 \times 10^{-8}$ | *Acadvl, Cpt2, Crot, Echs1, Etfb, Pex11g, Pex26, Stard7* |
| | DU6 | | |
| Lipid synthesis & storage | 76.5 | $3.8 \times 10^{-11}$ | *Hmgcr, Mogat2, Dagla, Daglb, Apobr, Ldlrap1, Stard5* |
